# Design and Biological Activity of a Novel Brain Penetrant Urea Compound Against Glioblastoma

**DOI:** 10.1101/2025.04.10.648292

**Authors:** Ling He, Xiaohong Chen, Hui Ding, Daria Azizad, Carter Hoffman, Linda Azizi, Gazmend Elezi, Zian Zhuang, Anjelica M. Cardenas, Carlos Calderon, Mahya Mohammadi, Sophia Tate, Julian Whitelegge, Gang Li, Jonathan E. Zuckerman, Robert Damoiseaux, Aparna Bhaduri, Linda M. Liau, Harley I. Kornblum, Michael E. Jung, Frank Pajonk

**Author notes:** **Correspondence address:** Michael E. Jung, PhD, Department of Chemistry at UCLA, 607 Charles E Young Dr E Los Angeles, CA 90095, Frank Pajonk, MD, PhD, Department of Radiation Oncology, David Geffen School of Medicine at UCLA, 10833 Le Conte Ave, Los Angeles, CA 90095-1714.

## Abstract

Glioblastoma (GBM) remains the most lethal primary brain tumor, largely due to therapy-resistant glioma stem cells (GSCs) and the ability of non-stem cells to dedifferentiate under therapeutic pressure. We developed MXC-017, a novel urea-based compound that crosses the blood–brain-barrier, directly targets GSCs, and prevents radiation-induced GSC formation. Using click chemistry pull-down and mass spectrometry, we identified vimentin as the target of MXC-017, further validated by *in silico* docking. Global transcriptomic profiling (bulk RNA-seq) and single-cell RNA-seq analyses revealed MXC-017’s efficacy with minimal off-target effects, supported by metabolic and kinome assays. Normal cell toxicity was negligible in fibroblasts, microglia, astrocytes, and murine neural progenitors. Maximum tolerated dose was identified and we observed significantly extended median survival in 17 PDOX GBM models when treated with MXC-017 plus radiation, benchmarked against standard-of-care temozolomide. These findings underscore the therapeutic potential of vimentin-targeting agents to overcome radiation resistance and improve outcomes for GBM patients.

**Statement of Significance:** Glioblastoma’s distinctive nature and the blood–brain barrier hamper therapies targeting therapy-resistant GSCs. We developed a novel urea-based agent that crosses the barrier, targets GSCs, and prevents radiation-induced GSC formation. With minimal off-target effects, reduced toxicity, and superior survival in PDOX models, it offers potential to improve outcome in GBM.

## Introduction

Despite advances in surgery and radiotherapy, glioblastoma (GBM) remains the deadliest adult brain cancer with unacceptable low median survival rates. Targeted therapies and biologics have failed to improve outcome and the standard-of-care against GBM has not changed in two decades.

GBM, like many solid tumors are thought to be organized hierarchically with a small number of radiation- and chemotherapy-resistant glioma stem cells (GSCs) producing more differentiated progeny and can repopulate tumors after sublethal treatment.

Several unique features of GBM add to the challenge of treating GBM and contribute to its inevitable recurrence. First, even though it rarely metastasizes outside of the CNS, GBM cells disperse into the normal parenchyma of the brain beyond the detectable tumor. Surgical access to these hidden cells is very limited and the maximum radiation dose to the whole brain is constrained by the normal tissue tolerance dose of the CNS. Even when the radiation dose to the detectable tumor is escalated, the median survival does not improve (1–3) and tumors often recur in close proximity to the initial tumors site and even within the radiotherapy target volume (4–6). Secondly, in animal models these dispersed cells have been shown to display an aggressive cancer stem cell phenotype even when compared to GSCs residing in the main tumor mass (7,8), thus suggesting that these cells are a major obstacle for controlling GBM with radiotherapy. Lastly, GBM cells show remarkable plasticity and non-stem glioma cells can dedifferentiate into treatment-induced GSCs (9–11) or transdifferentiate into endothelial- and pericyte-like cells to support tumor growth (12). Our recent data show that radiation induces a transient multipotent potent state in GBM cells that significantly enhances this intrinsic plasticity, leading to treatment resistance (9–12).

In this study we sought to develop novel compounds that cross the blood-brain barrier (BBB), target existing GSCs and interfere with glioma cell plasticity to prevent the induction of GSCs in response to irradiation. We show that novel urea compounds target existing GSCs, prevent the radiation-induced conversion of non-stem glioma cells into GSCs and prolong the median survival in mouse models of GBM with minimal normal tissue toxicity.

## Material & Methods

### Medicinal Chemistry

For the synthesis of MXC017, 4-Cyanophenylsulfonyl chloride **1** was converted into the sulfonamido primary amine **2** in two steps, formation of the sulfonamide and then hydride reduction of the nitrile. Reaction of the amine **2** with the indole phenyl carbamate **3**, prepared in one step from indole and diphenyl carbonate, in the presence of DBU gave the desired urea MXC017 in good yield (**Fig. 1E**). The other analogues were prepared by similar chemistry.

**Figure 1.**
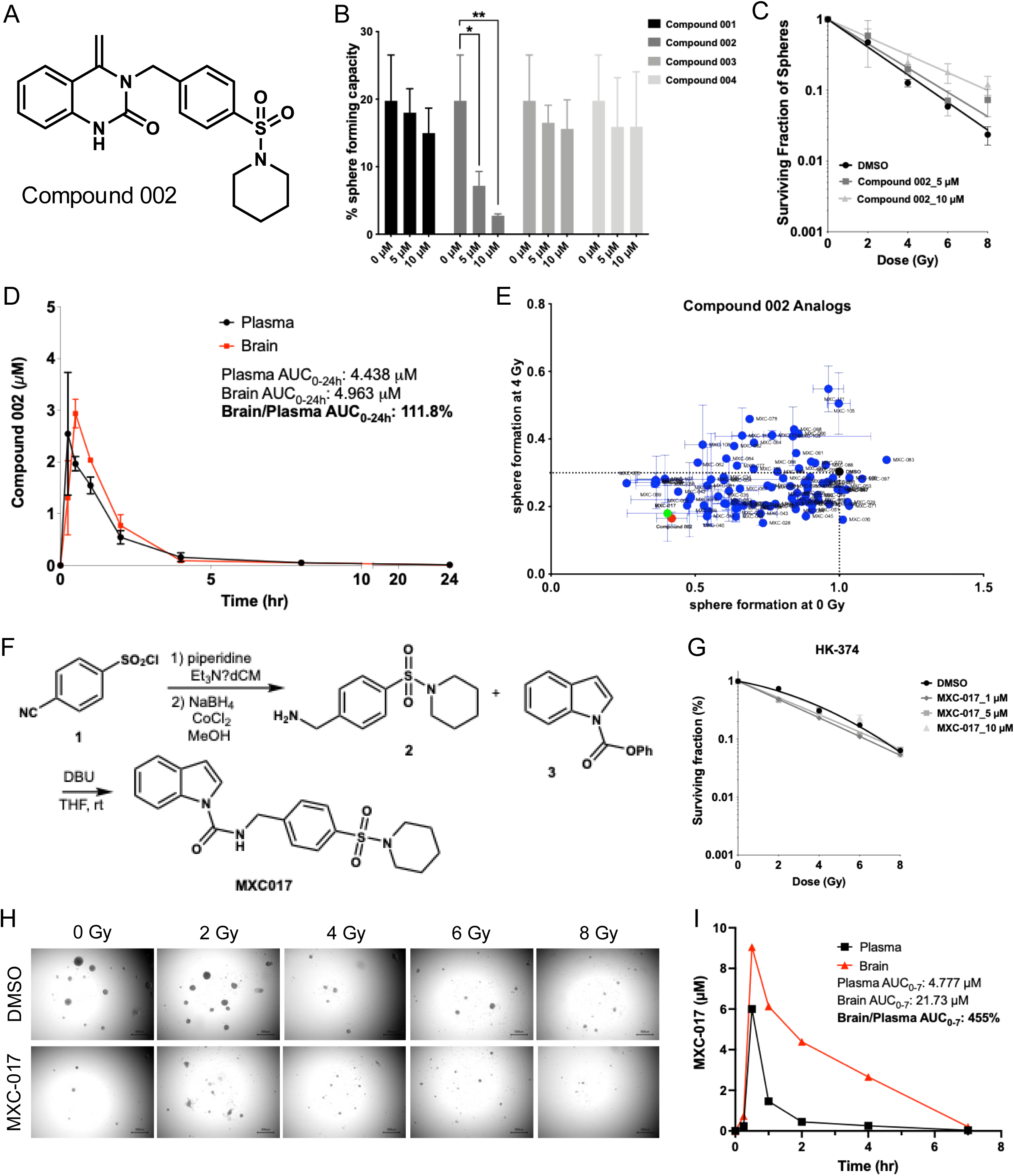
High-throughput screening and synthesis of lead compound MXC-017. **(A)** Chemical structure of compound 002. **(B)** Sphere-forming capacity of HK-374 patient-derived GBM cells treated with lead compounds 001-004 at concentrations of 5 and 10 µM. **(C)** Surviving fraction of HK-374 spheres treated with compound 002 (5 and 10 µM) following a single dose of 0, 2, 4, 6, or 8 Gy radiation. **(D)** Brain and plasma levels of compound 002 in C57BL/6 mice after a single intraperitoneal injection (50 mg/kg). **(E)** Sphere-forming capacity of HK-374 cells exposed to all 114 analogs of compound 002 under 0 Gy and 4 Gy conditions. **(F)** Synthesis scheme for the lead analog MXC-017. **(G)** Surviving fraction of HK-374 spheres treated with MXC-017 (1, 5, or 10 µM) following a single dose of 0, 2, 4, 6, or 8 Gy radiation. **(H)** Representative images of HK-374 spheres treated with 10 µM mxc-017 or DMSO (solvent control) following a single dose of 0, 2, 4, 6, or 8 Gy radiation. **(I)** Brain and plasma levels of MXC-017 in C57BL/6 mice after a single intraperitoneal injection (50 mg/kg). All experiments have been performed with at least 3 biological independent repeats. *P*-values were calculated using one-way ANOVA. * *p*-value < 0.05, ** *p*-value < 0.01.

### Cell lines

Primary human glioma cell lines were established at UCLA as described in (13); Characteristics of specific gliomasphere lines can be found in (14). Primary GBM cells were propagated as gliomaspheres in serum-free conditions in ultra-low adhesion plates in DMEM/F12, supplemented with SM1 Neuronal Supplement (#05177, STEMCELL Technology, Kent, WA), EGF (#78006, STEMCELL Technology), bFGF (#78003, STEMCELL Technology) and heparin (1,000 USP Units/mL, NDC0409-2720-31, Lake Forest, IL) as described previously (13–15).

For microglia cell cultures EOC 20 cells (female; sourced from C3H/HeJ mice) were obtained from ATCC (#CRL-2469, Manassas, VA). Cells were maintained under standard conditions in DMEM media/ 10 % FBS supplemented with 20 % conditioned media from CSF-1-expressing LADMAC bone marrow cells (#CRL-2420, ATCC). Normal human astrocytes (#CC-2565, Lonza) and NIH3T3 fibroblasts (#CRL-1685, ATCC) were cultured under standard conditions in DMEM supplemented with 10% FBS and 1% P/S. All cells were grown in a humidified atmosphere at 37°C with 5% CO_2_. The unique identity of all patient-derived specimens was confirmed by DNA fingerprinting (Laragen, Culver City, CA). All lines were routinely tested for mycoplasma infection (#G238, Applied biological Materials, Ferndale, WA).

### Irradiation

Cells were irradiated at RT using an experimental X-ray irradiator (Gulmay Medical Inc. Atlanta, GA) at a dose rate of 5.519 Gy/min. Control samples were sham-irradiated. The X-ray beam was operated at 300 kV and hardened using a 4 mm Be, a 3 mm Al, and a 1.5 mm Cu filter and calibrated using NIST-traceable dosimetry. Corresponding controls were sham irradiated.

For *in vivo* irradiation experiments, mice were anesthetized prior to irradiation with an intra-peritoneal injection of 30 µL of a ketamine (100 mg/mL, Phoenix, MO) and xylazine (20 mg/mL, AnaSed, IL) mixture (4:1) and placed on their sides into an irradiation jig that allows for irradiation of the midbrain while shielding the esophagus, eyes, and the rest of the body.

### Drug treatment

For *in vitro* studies, MXC-017 was dissolved at a stock concentration of 10 mM in DMSO, and 1 µM, 5 µM, 10 µM were used. Temozolomide was prepared at a stock concentration of 10 mM in DMSO, 1 µM, 3 µM, 6 µM, and 9 µM were used.

For the *in vivo* survival studies, tumor engraftment was confirmed using bioluminescence imaging. Mice were then treated intraperitoneally with MXC-017 at either 50 mg/kg or 150 mg/kg on a 5-days on / 2-days off schedule or were given temozolomide at 66 mg/kg by oral gavage for two weeks, until they reached predefined euthanasia endpoints. MXC-017 was dissolved in corn oil containing 2.5% DMSO at a concentration of 5.5 mg/ml and prepared freshly for the injections. Likewise, temozolomide was freshly prepared at 6 mg/ml in corn oil with 2.5% DMSO before administration.

*In-vitro* limiting dilution assay

For the assessment of self-renewal *in vitro*, primary GBM cells were irradiated with 0, 2, 4, 6 or 8 Gy and seeded under serum-free conditions into non-tissue-culture-treated 96-well plates in DMEM/F12 media, supplemented with 10 mL / 500 mL of B27 (Invitrogen), 0.145 U/mL recombinant insulin (Eli Lilly, Indiana), 0.68 U/mL heparin (Fresenius Kabi, Illinois), 20 ng/mL fibroblast growth factor 2 (bFGF, Sigma) and 20 ng/mL epidermal growth factor (EGF, Sigma). The number of cells seeded per well was optimized for each GBM cell line through seeding a serial dilution of cells across different wells. The number of spheres formed at each dose point was normalized against the non-irradiated control. The resulting data points were fitted using a linear-quadratic model.

For Secondary and tertiary glioblastoma spheres, primary GBM spheres were grown in 10-cm petri dishes and treated with 10 µM MXC-017one hour before a single dose of 4 Gy radiation. The irradiated spheres were dissociated after 5-7 days and plated at clonal densities to generate secondary spheres. Re-plated secondary spheres were treated with a single dose of 4 Gy, with or without MXC-017 (10 µM). 5-7 days later, the spheres were dissociated and plated at clonal densities to form tertiary spheres.

### Mass Spectrometry for Pharmacokinetics study

Sample Preparation: Whole blood from mice was centrifuged to isolate plasma. MXC-017 was isolated by liquid-liquid extraction from plasma: 50 µl plasma was added to 2 µl internal standard and 100 µl acetonitrile. Mouse brain tissue was washed with 2 mL cold saline and homogenized using a tissue homogenizer with fresh 2 mL cold saline. MXC-017 was then isolated and reconstituted in a similar manner by liquid-liquid extraction: 100 µl brain homogenate was added to 2 µl internal standard and 200 µl acetonitrile. The samples were centrifuged, supernatant removed and evaporated by a rotary evaporator and reconstituted in 100 µl 50:50 water:acetonitrile.

MXC-Detection: Chromatographic separations were performed on a 100 x 2.1 mm Phenomenex Kinetex C18 column (Kinetex) using the 1290 Infinity LC system (Agilent). The mobile phase was composed of solvent A: 0.1% formic acid in Milli-Q water, and B: 0.1% formic acid in acetonitrile. Analytes were eluted with a gradient of 5% B (0-4 min), 5-99% B (4-32 min), 99% B (32-36 min), and then returned to 5% B for 12 min to re-equilibrate between injections. Injections of 20 µl into the chromatographic system were used with a solvent flow rate of 0.10 ml/min.

Mass spectrometry was performed on the 6460 triple quadrupole LC/MS system (Agilent). Ionization was achieved by using electrospray in the positive mode and data acquisition was made in multiple reactions monitoring mode. For MXC-017: m/z 384.2→ 253.1 with a fragmentor voltage of 140V, and collision energy of 16 and 20 eV, respectively. The analyte signal was normalized to the internal standard and concentrations were determined by comparison to the calibration curve (0.5, 5, 50, 250, 500, 2000 nM). MXC-017 brain concentrations were adjusted by 1.4% of the mouse brain weight for the residual blood in the brain vasculature as described by Dai *et al* (16).

### Reprogramming Assay

HK-374 ZsG-cODC cells were cultured as monolayers, and ZsG-negative cells were sorted using a BD FACSAria™ III sorter (BD Biosciences). Sorted cells were plated in 6-well plates at a density of 50K per well. The next day, cells were treated with 10 µM MXC-017, followed by irradiation at 0 or 4 Gy after 1 hour. Plates were incubated for 5 days post-irradiation, then cells were analyzed for green fluorescence via flow cytometry using an LSRFortessa™ Cell Analyzer (BD Biosciences). Non-ZsG-transfected cells served as controls for gating. Data were processed with FlowJo software, and fold increases in ZsG-positive cells were calculated relative to the 0 Gy control.

### Extreme Limiting Dilution Analysis (ELDA)

3×10^5^ HK-374 cells were intracranially implanted into the NSG mice as described above. Tumors were grown for 3 days for successful grafting. Tumor-bearing mice were then treated intra-peritoneally with either corn oil or MXC-017 and irradiated with a single dose of 4 Gy one hour after the next day. The MXC-017 injection was on a 5-days on / 2-days off schedule for 2 weeks. The mice were then euthanized and tumor-bearing brains were dissected and further subjected for dissociation using mouse Tumor Dissociation Kit (# 130-096-730, Miltenyi, Auburn, CA) to get single cell suspension, as described in (10). The cells were counted and plated into the non-tissue-culture-treated 96-well plates at a range of 1 to 512 cells/well. Growth factors (EGF and bFGF) were supplemented every two days. Glioma spheres were counted 10 days later and presented as the percentage to the initial number of cells plated. The glioma stem cell frequency was calculated using the ELDA software (17).

### Cell cycle analysis

Following treatment with 10 µM MXC-017, HK-374 cells were trypsinized, rinsed with ice-cold PBS, and fixed in cold 70% ethanol while gently vortexing. The cells were centrifuged at 400 *x g* for 4 minutes, resuspended in 200 µl UltraPure RNase (Thermo Fisher, #12-091-021), and transferred to FACS tubes. Next, 200 µl of Propidium Iodide solution (1 mg/ml, Thermo Fisher, #P1304MP) was added, and the cells were incubated for 15 minutes at room temperature in the dark. Flow cytometry was performed on at least 100,000 events using an LSR Fortessa (BD Biosciences, Franklin Lakes, NJ).), and data were analyzed with FlowJo v10.

### Annexin V/PI Staining and Flow Cytometry

HK-374 monolayer cells were plated onto 10-cm petri dishes and treated with MXC-017 (10 µM) one hour prior to receiving a single dose of 4 Gy irradiation the following day. 72 hours post-irradiation, cells were harvested, washed twice with cold PBS, and resuspended in 1X Annexin V binding buffer at a concentration of 1 × 10 cells/ml. For each sample, 100 μl of cell suspension was incubated with 5 μl of FITC-conjugated Annexin V and 5 μl of propidium iodide (PI, 50 μg/ml) for 15 minutes at room temperature in the dark. Following incubation, 400 μl of binding buffer was added, and samples were immediately analyzed using a BD LSR Fortessa flow cytometer (BD Bioscience, San Jose, CA), acquiring a minimum of 100,000 events per sample. Data analysis were performed using FlowJo v10 software. Cells positive for Annexin V and negative for PI were defined as early apoptotic, double-positive cells as late apoptotic or necrotic, and double-negative cells are viable cells.

### Animals

Both female and male 6–8-week-old NOD-*scid* IL2Rgamma^null^ (NSG), originally obtained from The Jackson Laboratories (Bar Harbor, ME) were re-derived, bred and maintained in a pathogen-free environment in the American Association of Laboratory Animal Care-accredited Animal Facilities of Department of Radiation Oncology, University of California, Los Angeles. Patient-derived GBM cells (2×10^5^ cells, a cell line list is provided in the **Suppl. Table 1**) were implanted into the right striatum of the brains of mice using a stereotactic frame (Kopf Instruments, Tujunga, CA) and a nano-injector pump (Stoelting, Wood Dale, IL). The injection coordinates were 0.5 mm anterior and 2.25 mm lateral to the bregma, at a depth of 3.0 mm from the brain surface. Tumors were allowed to grow for 3-7 days, depending on the growth rate, with successful grafting confirmed by bioluminescence imaging. Mice that lost 20% of their body weight or developed neurological deficits necessitating euthanasia were sacrificed.

### Click-Chemistry affinity capture of protein targets

To generate MXC-017 affinity beads, alkyne-tagged MXC-017 was synthesized and coupled to azide agarose beads (Sigma, #900957). Briefly, azide agarose beads were incubated with alkyne-tagged MXC-017 at a final concentration of 5 mM in coupling buffer (PBS containing 20% DMSO) for 4 hours at room temperature. After coupling, the beads were washed with PBS to remove unbound ligand, then blocked with 1 M Tris-HCl (pH 7.5) for an additional hour. Control beads were prepared by blocking uncoupled azide agarose beads with 1 M Tris-HCl (pH 7.5).

To identify proteins interacting with MXC-017, 6 x 10^7^ HK-374 cells were homogenized in 2 mL cell lysis buffer (50 mM Tris-HCl, pH 7.5, 150 mM NaCl, 2 mM EDTA, 0.5% NP-40) on ice for 30 min. The homogenate was then centrifugated at 12,000 x g for 15 min. The supernatant was divided into 1 mL aliquots and mixed with 100 µl of either control beads or MXC-017 agarose beads in the presence of 2% DMSO or 1.25 mM MXC-017, followed by rotation for 1 hour at RT. The beads were collected by centrifugation and washed three times with 1 mL of cell lysis buffer each. The beads were then boiled in SDS sample buffer, centrifuged, and the supernatant was run on a 10% SDS-PAGE gel, followed by staining with Coomassie Blue. Following the de-stain procedure, the targeted bands were excised, trypsin-digested and subjected to mass spectrometry for protein identification.

### Mass Spectrometry for Click-Chemistry pulldown

#### In-Gel digestion and extraction

Gel bands were destained in 200 uL of 100 mM ammonium bicarbonate/50% ACN at 37°C for 10 min and were repeated for three times. Gel bands were dried in speed vac for 5 min before addition of 50 μL of 0.01 mg/mL trypsin solution and incubation at 37°C overnight. Protein digests were extracted by 50 μL of 50% ACN/0.1% formic acid for three times. Gel extracts were combined for desalting by Empore stage-tip.1 Elution from the stage-tip was dried by speed vac and re-suspended in 10.0 uL 3% acetonitrile with 0.1% formic acid.

#### LC MS/MS

1.0 uL sample was injected to an ultimate 3000 nano LC, which was equipped with a 75 µm x 2 cm trap column packed with C18 3 µm bulk resins (Acclaim PepMap 100, Thermo Scientific) and a 75 µm x 15 cm analytical column with C18 2 µm resins (Acclaim PepMap RSLC, Thermo Scientific). The nanoLC gradient was 3−35% solvent B (A = H2O with 0.1% formic acid; B = acetonitrile with 0.1% formic acid) over 40 minutes and from 35% to 85% solvent B in 5 min at flow rate 300 nL/min. The nannoLC was coupled with a Q Exactive orbitrap mass spectrometer (Thermo Fisher Scientific, San Jose, CA). The ESI voltage was set at 1.9 kV, and the capillary temperature was set at 275°C. Full spectra (m/z 350 - 2000) were acquired in profile mode with resolution 70,000 at m/z 200 with an automated gain control (AGC) target of 3 × 10^6^. The most abundance 15 ions were subjected to fragmentation by higher-energy collisional dissociation (HCD) with normalized collisional energy of 25. MS/MS spectra were acquired in centroid mode with resolution 17,500 at m/z 200. The AGC target for fragment ions are set at 2 × 10^4^ with maximum injection time of 50 ms. Charge states 1, 7, 8, and unassigned were excluded from tandem MS experiments. Dynamic exclusion was set at 45.0 s.

#### Data Analysis

Raw data was searched again uniprot human database by Proteome Discoverer version 2.5. Following parameters were set: precursor mass tolerance ± 10 ppm, fragment mass tolerance ± 0.02 Th for HCD, up to two miscleavages by trypsin, methionine oxidation as variable modification.

### Immunofluorescent staining

For in vivo tumor specimen, brain sections were baked for 30 min in an oven at 65 °C, deparaffinized in two successive Xylene baths for 5 minutes each and then hydrated for 5 minutes each using an alcohol gradient (ethanol 100%, 90%, 70%, 50%, 25%). Antigen retrieval was performed using Heat Induced Epitope Retrieval in a citrate buffer (10 mM sodium citrate, 0.05% tween20, pH 6) with heating to 95 °C in a steamer for 20 minutes. After cooling down, the slides were blocked with 10% goat serum plus 1% BSA at room temperature for 30 minutes and then incubated with the primary antibodies against olig2 (#65915s, cell signaling technology, 1:400) mixed with nestin (#33475s, cell signaling technology, 1:2000) or GFAP (#80788s, cell signaling technology, 1:400) mixed with human mitochondria (ab92824, Abcam, 1:800) overnight at 4°C. The secondary antibodies Alexa Fluor 594 Goat Anti-rabbit immunoglobulin G (IgG) (H/L) antibody (1:1,000 (Invitrogen)) and Alexa Fluor 488 Goat Anti-mouse IgG (H/L) antibody (1:1,000 (Invitrogen)) were applied followed by nuclear counterstaining and mounting procedures as above. Fluorescent images were taken with a digital microscope (BZ-9000, Keyence, Itasca, IL).

For *in vitro* vimentin study, HK-374 monolayers were plated onto the 2-chamber cell culture slides (#154461, ThermoFisher Science) and treated with 10 µM MXC-017 with or without a single dose of 4 Gy irradiation. 6 hours and 24 hours post-irradiation, cells were fixed, permeabilized, blocked and stained with monoclonal vimentin antibody (#5741s, cell signaling technology, 1:100) or polyclonal vimentin antibody (ab137321, Abcam, 1:500), followed by Alexa Fluor 488 Goat Anti-rabbit IgG (H/L) secondary antibody (1:1,000 (Invitrogen)) as described above. Fluorescent images were then acquired using a confocal microscope (Zeiss).

### Western Blotting

HK-374 cells were serum-starved overnight to synchronize cellular signaling. The following day, cells were treated with 10 µM MXC-017 or DMSO (vehicle control), and whole cell lysates were collected at 15 minutes, 30 minutes, 1 hour, 2 hours, 4 hours, and 6 hours post-treatment. In a separate experiment, cells were treated with 10 µM MXC-017 or DMSO for 2 hours, after which whole cell lysates were collected. The culture medium was then replaced with complete medium containing serum and freshly prepared 10 µM MXC-017, and cells were incubated for an additional 30 minutes before harvesting lysates. Following this, the medium was switched back to serum-free conditions with another round of MXC-017 treatment. Additional lysates were collected at 30 minutes and 2 hours after this final treatment.

To prepare whole cell lysates, cells were lysed in RIPA lysis buffer containing proteinase inhibitor and phosphatase inhibitor. 150 µl of ice-cold RIPA lysis buffer (10 mM Tris-HCl (pH 8.0), 1 mM EDTA, 1% Triton X-100, 0.1% Sodium Deoxycholate, 0.1% SDS, 140 mM NaCl, 1 mM PMSF) containing proteinase inhibitor (Thermo Fisher Scientific) and phosphatase inhibitor (Thermo Fisher Scientific) was used to lyse the cells. The protein concentration in each sample was determined by BCA protein assay (Thermo Fisher Scientific) and samples were denaturated in 4x Laemmli sample buffer (Bio-Rad) containing 10% β-mercaptoethanol for 10 minutes at 95 °C. Equal amounts of protein were loaded onto 10% SDS-PAGE gels and subjected to electrophoresis for 2 hours. Samples were then transferred onto 0.45 µm nitrocellulose membrane (Bio-Rad) and blocked in 1x TBST containing 5% bovine serum albumin (BSA) for 30 minutes at RT, followed by incubation with primary antibodies against vimentin (#ab137321, 1:1000, Abcam), Phospho-vimentin (Ser83) (#3878S, 1:1000, Cell Signaling Technology), Phospho-vimentin (Ser39) (#13614S, 1:1000, Cell Signaling Technology), Phospho-vimentin (Ser56) (#3877S, 1:1000, Cell Signaling Technology), and β-actin (#3700S, 1:1000, Cell Signaling Technology) in 1X TBST containing 5% BSA overnight at 4°C with gentle rocking. Membranes were then washed three times for 5 minutes each with 1X TBST and incubated with secondary antibodies, 1:5000 anti-mouse or anti-rabbit horseradish peroxidase (HRP; Cell Signaling) in 1X TBST for two hours at RT with gentle rocking. Membranes were washed again three times for 5 minutes each with 1X TBST. Pierce ECL Plus Western Blotting Substrate (Thermo Fisher) was added to each membrane and incubated at RT for 5 minutes. The blots were then scanned using the Odyssey Fc imaging system (LI-COR Biosciences, Lincoln, NE). β-actin served as a loading control.

### *In Silico* Docking Analysis

*In silico* docking was performed using the MolModa software package vers. 1.0.1 (18). A truncated tetramer model of vimentin (PDB ID: 3KLT) and a complete atomic model of vimentin (PDB ID: 8RVE) (19) were obtained from from the PDB data base. Potential binding pochets were identified using FPocketWeb vs 1.0.1. Docking was performed using the AutoDock Vina algorithm (20) and illustrated using ChimeraX vers. 1.9.

### Migration/Invasion Assay

Serum-starved HK-374 monolayers were trypsinized and plated onto Transwell inserts (#08-771-12, Fisher Scientific) at a density of 1 x 10^5^ cells per well. MXC-017 or solvent control (DMSO) was added to the medium in the bottom chamber containing 10% FBS. A negative control using 1% FBS medium was included to eliminate the FBS gradient between the upper and lower chambers. After 16-24 hours of incubation, cells were fixed with 10% formalin. Non-migrated cells on the upper side of the membrane were gently removed with cotton swabs, while migrated cells on the lower side were stained with 0.5% crystal violet. Images were captured using a digital microscope (BZ-9000, Keyence, Itasca, IL) and quantified using Image J software.

For experiments involving irradiation, HK-374 monolayers were plated onto 10-cm Petri dishes and treated with 10 µM MXC-017, with or without a single dose of 4 Gy irradiation. 48 hours post-irradiation, the cells were serum starved overnight. The following day, cells were trypsinized and plated onto Transwell inserts at 1 x 10^5^ cells per well. The bottom chamber contained 10% FBS medium to create an FBS gradient that promotes cell migration. After 16-24 hours, cells were fixed and stained with crystal violet as described above.

### Metabolomic assay

HK-374 monolayers were plated onto 6-well plates at a density of 100K cells per well for 0 Gy conditions, while 150K cells per well for 4 Gy conditions. Cells were incubated with complete media containing 17.5 mM [1,2-^13^C2]-glucose (99%) (Cambridge Isotope Laboratories), and treated with MXC-017 at 10 µM one hour before a single dose of 4 Gy irradiation. Metabolites were extracted at 48 hours post-irradiation as previously described, with 80% methanol 0.1 M formic acid at -80°C, and neutralized with 15% (w/v) ammonium bicarbonate.

The metabolites were loaded onto a Luna 3um NH2 100A 150 × 2.0 mm column (Phenomenex). The chromatographic separation was performed on a Vanquish Flex (Thermo Scientific) with mobile phases A (5 mM ammonium acetate buffer at pH 9.9) and B (acetonitrile) and a flow rate of 200 μL/min. A linear gradient from 15% A to 95% A over 18 minutes was followed by 9 min isocratic flow at 95% A and re-equilibration to 15% A. Metabolites were detected with a Thermo Scientific Q Exactive mass spectrometer run with polarity switching (+3.5 kV/− 3.5 kV) in full scan mode with an m/z range of 70-975. TraceFinder 4.1 (Thermo Scientific) was used to quantify metabolites by area under the curve using expected retention time and accurate mass measurements (< 5 ppm). The 13C labeling measurements from LC-MS were corrected to account for natural 13C isotope abundance and ^12^C impurity present in the ^13^C-glucose tracers.

### Whole kinome screening

The KINOMEscan™ platform, offered by Eurofins DiscoverX, is a high-throughput kinase profiling assay designed to measure the binding affinity and selectivity of small-molecule inhibitors against a broad panel of kinases. For this assay, a 10 µM solution of MXC-017, prepared in DMSO, was sent to Eurofins. The Full Kinase Profiler Panel [Km ATP] (#50-000KP, Eurofins) was utilized for the analysis.

A Kinase HotSpot assay was conducted to evaluate the inhibitory potential of Compound MXC-017 against the selected kinase targets FLT3, JNK3, and TNIK. A 10 µM solution of MXC-017, prepared in DMSO, was shipped to Reaction Biology Corporation (Malvern, PA) for profiling. Each kinase was tested using an ATP concentration set to its respective Michaelis constant (Km ATP). After the initial screening, the half-maximal inhibitory concentration (IC_50_) values were determined to quantify MXC-017’s potency against these specific enzymes.

### Bulk RNA sequencing

Primary HK-374 cells were treated with 10 µM MXC-017 one hour before a single dose of radiation (4 Gy), RNA was extracted at 48 hours using Trizol followed with RNeasy kit (Qiagen) isolation. The next-generation sequencing was performed by Novogene (Chula Vista, CA) and reads were mapped to the human genome (hg38) following their standard pipeline (10). Read counts were analyzed using the iDEP package (version 2.0) (21). Differentially expressed genes were calculated using the DESeq2 algorithm with a minimum of a 1.5-fold change and a false discovery rate (FDR) of 0.1. Enrichment *p*-values were calculated based on a one-sided hypergeometric test. *P*-values were then adjusted for multiple testing using the Benjamini-Hochberg procedure and converted to FDR. Fold Enrichment was defined as the percentage of genes in the list belonging to a pathway, divided by the corresponding percentage in the background. Overlap of differential expressed genes with known stem cell signatures was calculated using the StemChecker tool (22) with both "Cell Proliferation Genes" and "Cell Cycle Genes" masked.

### Single Cell RNA sequencing

HK-374, HK-390, HK-217, and HK-244 gliomaspheres were plated onto the Poly-D-Lysine/Laminin coated 6-well plate (#354595, Corning) and treated with MXC-017 at a concentration of 10 µM, one hour before receiving a single dose of 4 Gy the next day. Five days post-irradiation, the gliomaspheres in suspension culture were collected and dissociated using TrypLE (no phenol red, Thermo Fisher Scientific). The adherent differentiated cells were washed with HBSS and de-attached using Trypsin/EDTA (Cell Applications, Cat#090K). The cells were then pooled, filtered through a 40-µm strainer and counted with a hemacytometer. The samples were pelleted by centrifugation at 300 g for 5 minutes at 4°C, resuspended in the provided Cell Resuspension Buffer (Fluent Bioscience, Watertown, MA), and 50,000 cells were targeted for capture and processed for single-cell transcriptomics using the PIPseq^TM^ T2 3’ Single Cell RNA Kit v4.0 Plus and v5 (Fluent Biosciences).

For sequencing of *in vivo* samples, 300,000 HK-374 cells, engineered to express GFP and firefly luciferase, were implanted intracranially and allowed to engraft for one week. Engraftment was verified by bioluminescent imaging, after which mice received either a single 4 Gy dose of radiation alone or radiation followed by MXC-017 treatment (5 i.p. injections per week, 50 mg/kg) for two weeks. Tumors were then harvested and dissociated into single cells using the Miltenyi Brain Tumor Dissociation Kit (#130-095-942), followed by mouse cell depletion (Mouse Cell Depletion Kit, #130-104-694, Miltenyi). Cells from 2-3 mice were pooled, passed through a 40-µm strainer and counted using a hemacytometer. After pelleting by centrifugation at 300 g for 5 minutes at 4°C, the cells were resuspended in the provided Cell Resuspension Buffer. Approximately 50,000 cells were then captured and processed for single-cell transcriptomics using the PIPseq^TM^ T2 3’ Single Cell RNA Kit v5 (Fluent Biosciences).

Sequencing libraries were prepared according to PIPseq T2 instructions, with 12 cycles for cDNA amplification and 8 cycles for library amplification, depending on the DNA amount. The libraries were quantified using a Qubit 4 Fluorometer and a TapeStation 4200 and sequenced as per manufacturer recommendation on a NovaSeq X Plus – 25B 2 x150bp, with 3.25 billion reads in total for 8 samples.

### scRNA sequencing data analysis

FASTQ files were aligned to the human genome (GRCh38) and processed using the PIPseeker software, version 3.3.3 (Fluent Biosciences). Raw matrices were in processed in Seurat version 5 R package (23) in RStudio. Cells with less than 500 genes were filtered out and cells with high mitochondrial gene count (>5%) were removed. Downstream analyses and annotations were performed as described in (24). Senescence indices were calculated using the SenePy Python package (25).

### Clonogenic survival assay

EOC20, Normal human astrocytes, and NIH3T3 fibroblasts were trypsinized and plated in 6-well plates at a density of 100-400 cells. The cells were treated with C2 (5 µM), or MXC-017 (1, 2.5, 5, and 10 µM) the next day, DMSO served as solvent control. After two weeks, the colonies were fixed and stained with 0.5 % crystal violet. Colonies consisting of at least 50 cells were counted in each group and presented as percentage of the initial number of cells plated.

### Maximum Tolerated Dose

6-8 weeks male and female C57BL/6 mice were used. Animals were treated in pairs (1 male/1 female) with a single dose of MXC-017 at 150, 300, 600, and 900 mg/kg administered intraperitoneally (i.p.). The mice were observed and weighted for two weeks. After euthanasia all external body orifices were examined, blood was drawn via cardiac puncture, and hematological parameters were analyzed using HemaVet^®^950FS (The Drew Company, Boston, MA). Body cavities (abdomen, thorax, and cranium) were opened and inspected, and major organs (brain, lung, heart, liver, kidneys, and spleen) were subjected to histopathological examination by a board-certified pathologist.

For the study of repeated compound administration, mice were treated in groups of 6 mice (3 male/3 female) at doses of 150, 300, and 600 mg/kg via i.p. for 5 consecutive days per week over 2 weeks. The animals were weighted daily and monitored for clinical signs of MXC-017 toxicity. Two days after the final injection, the mice were subjected to terminal necropsy, and their bodies, blood, and major organs were examined as described above.

### Clinical Chemistry

6-8 weeks male and female C57BL/6 mice were administered MXC-017 at 150 or 300 mg/kg intraperitoneally for five consecutive days each week over a two-week period. Serum samples were then collected and tested using the Comprehensive Plus Clinical Chemistry Panel (#80824, VRL, Maryland).

### Ethics statement

All animal experiments were approved by UCLA’s Institutional Animal Care and Use Committee in accordance with all local and national guidelines for the care of animals.

### Statistics

All data shown are represented as mean ± standard error mean (SEM) of at least 3 biologically independent experiments. A *p*-value of ≤0.05 in an unpaired two-sided *t*-test or one-sided ANOVA for multiple testing indicated a statistically significant difference. Kaplan-Meier estimates were calculated using GraphPad Prism Software (version 10.2.0). A *p*-value of 0.05 or less in the log-rank test indicated a statistically significant difference.

## Results

### High-throughput screen and synthesis of the lead compound MXC-017

In a recent high-throughput screen (HTS) of over 83,000 compounds including FDA-approved compounds we identified potential inhibitors of radiation-induced tumor cell plasticity (26). Restricting our selection to compounds with an at least a two-fold reduction in radiation-induced phenotype conversion of non-stem tumor cells into cancer stem cells, no Lipinski violations (27), a CNS multiparameter optimization score (28) of 4 or higher, and no radiosensitizing effect, we identified compound 002 (4-methylidene-3-[[4-(piperidine-1-sulfonyl)phenyl]methyl]-1,2,3,4-tetrahydroquinazolin-2-one, Molport-009-476-782) as a potential lead candidate **(Fig. 1A)**. This compound significantly reduced self-renewal capacity in HK-374 patient-derived GBM cells without inducing radiosensitization **(Fig. 1B/C)**. Mass spectrometry of plasma and brain tissue of non-tumor bearing mice confirmed that compound 002 crossed the BBB (**Fig. 1D**). However, when we synthesized compound 002, it lacked the anti-tumor properties of compound 002 obtained from Molport. Subsequent NMR spectroscopy of Molport-009-476-782 revealed several degradation products **(Suppl. Fig 1)**, which led us to synthesize 114 novel analogs of compound 002 and to test them for their ability to prevent spontaneous (0 Gy) and radiation-induced (4 Gy) phenotype conversion of non-stem GBM cells into GSCs **(Fig. 1E)**. Structures of all 114 compounds can be found in the **Suppl. Table 2**. Among these, MXC-017 emerged as the lead compound. A description of the synthesis process is shown in **Fig. 1F**. Additionally, MXC-017 displayed no radiosensitizing effect yet caused a significant reduction in sphere formation **(Fig. 1G/H)**. Mass spectrometry of plasma and brain tissue demonstrated the superior BBB penetration of MXC-017 (brain-to-plasma ratio of 4.55; **Fig. 1I**), surpassing even the psychotropic drug quetiapine (brain-to-plasma ratio of 3.08) (9)

### MXC-017 inhibits cellular plasticity in GBM and prolongs survival in a PDOX GBM mouse model

Our HTS was performed using our imaging system for putative GSCs with low proteasome activity (15). Using this system we confirmed radiation-induced induction of marker-positive cells and its prevention by treatment with MXC-017 in HK-374 cells (classical TCGA subtype; **Fig. 2A**). The effect of MXC-017 on sphere-forming capacity, a functional measure of self-renewal, was dose-dependent and comparable to that of the original compound 002 **(Fig. 2B)**. Likewise, MXC-017 significantly reduced sphere-formation in HK-345 (mesenchymal TCGA subtype; **Fig. 2C**) and to a lesser extend in HK-157 cells (proneural TCGA subtype; **Fig 2D**).

**Figure 2.**
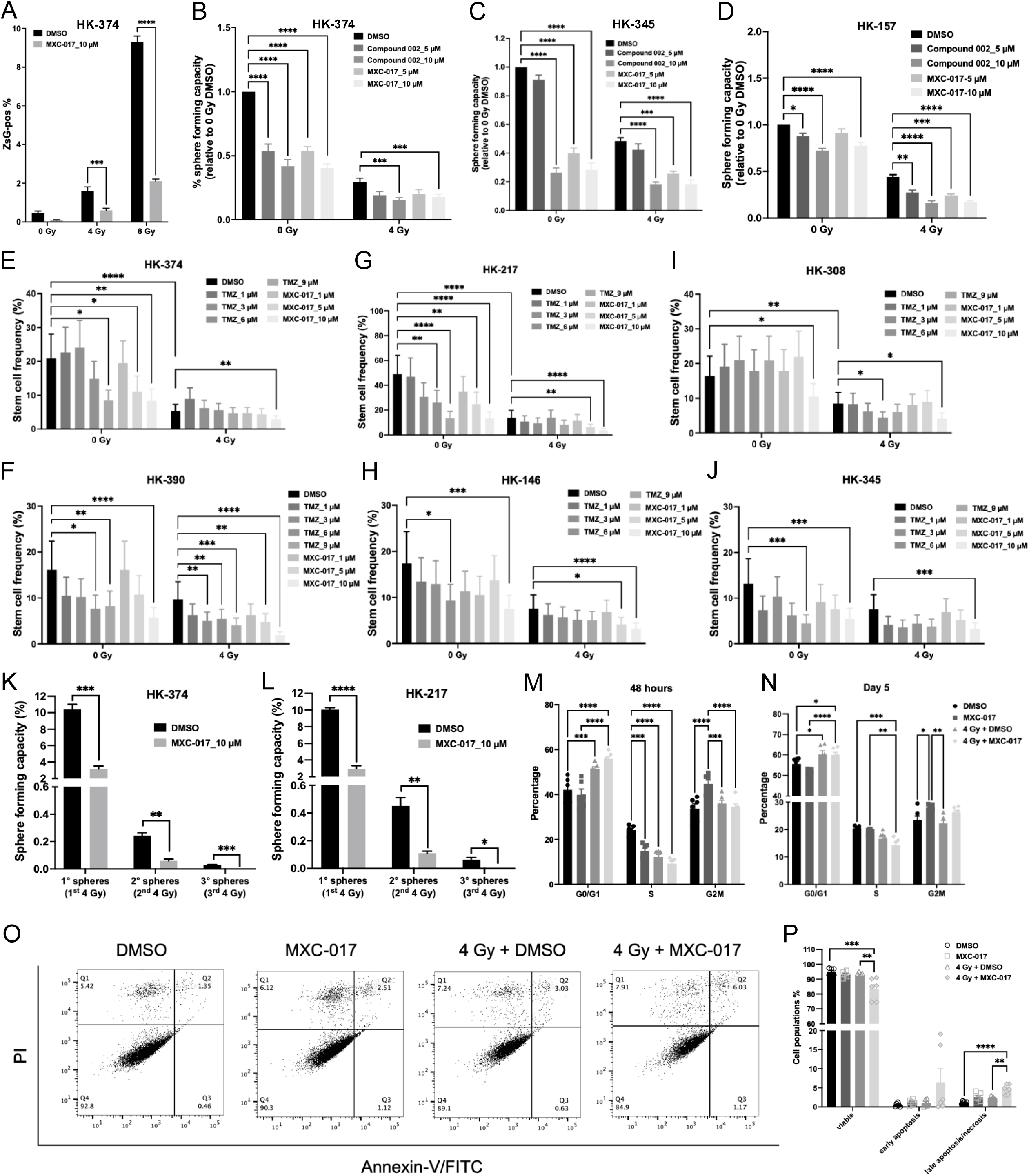
MXC-017 inhibits cellular plasticity in GBM. **(A)** Sorted ZsGreen-cODC-negative HK-374 ZsGreen-cODC vector expressing cells were plated (50,000 cells/well in 6-well plates) and pre-treated with MXC-017 (10 µM) or DMSO for one hour before receiving a single dose of 0, 4,or 8 Gy radiation. Five days later, cells were trypsinized and analyzed for ZsGreen-cODC-positive population via flow cytometry, with non-infected parental HK-374 cells served as controls. **(B-D)** Sphere-forming capacity of HK-374, HK-345, and HK-157 GBM cells treated with MXC-017 (5 or 10 µM) after a single dose of 0 or 4 Gy radiation, compared to compound 002 (5 or 10 µM). **(E-J)** Stem cell frequency of HK-374 and HK-390 (classical), HK-217 and HK-146 (proneural), HK-308 and HK-345 (mesenchymal) GBM cells treated with MXC-017 (1, 5, or 10 µM) following a single dose of 4 Gy radiation, benchmarked against standard-of-care temozolomide (1, 3, 6, or 9 µM) plus 4 Gy radiation. **(K/L)** Sphere-forming capacity of first, secondary, and tertiary HK-374 and HK-217 spheres treated with MXC-017 combined with a single dose of 4 Gy radiation. **(M/N)** Cell cycle distribution of HK-374 cells treated with MXC-017 (10 µM) in the presence or absence of a single dose of 4 Gy at 48 hours and day 5. **(O)** Representative Annexin V/PI dot plots of HK-374 cells treated with MXC-017 (10 µM), with or without a single 4 Gy dose of radiation, analyzed 72 hours post-treatment. **(P)** Quantification of viable, early apoptotic, and late apoptotic/necrotic cell populations. All experiments have been performed with at least 3 biological independent repeats. *P*-values were calculated using unpaired t-test for A, K, L; one-way ANOVA for B, C, D, M, N, P; two-way ANOVA for E-J. * *p*-value < 0.05, ** *p*-value < 0.01, *** *p*-value < 0.001, **** *p*-value < 0.0001.

To test the effect of MXC-017 on GSC frequencies we performed ELDAs using HK-374 and HK-390 cells (both classical TCGA subtype; **Fig. 2E/F**), HK-217 and HK-146 cells (both proneural TCGA subtype; **Fig. 2G/H)**, and HK-308 and HK-345 cells (both mesenchymal TCGA subtype; **Fig. 2I/J**) and compared it to the effect of temozolomide (TMZ) at concentrations reported for glioma tissue in human patients under TMZ treatment(29). In all six lines, MXC-017 outperformed TMZ and significantly reduced GSC frequencies alone or in combination with radiation **(Suppl. Tables 3 and 4)**. To exclude that MXC-017 would eliminate only a subpopulation of GSCs and select for resistant subclones, we performed secondary and tertiary clonal sphere-forming assay using HK-374 and HK-217 cells and confirmed that MXC-017 accelerated GSC exhaustion in both lines (**Fig. 2K/L**). Cell cycle analysis showed that MXC-017 enhanced the radiation-induced G_0_/G_1_ arrest and reduced the number of cells in S-phase, at 48 h and 5 days after irradiation (**Fig. 2M/N**). FACS analysis using Annexin V/PI staining showed that irradiation with 4 Gy or treatment with MXC-17 alone did not induce apoptosis. However, combined treatment with radiation and MXC-017 significantly reduced the viable cell population and increased the proportion of apoptotic cells at 72 hours post-treatment (**Fig. 2O/P**).

To test if MXC-017 would affect GSCs *in vivo,* HK-374 cells were orthotopically injected into the brains of NSG mice. Grafting was confirmed by BLI and mice were irradiated with a single dose of 4 Gy, treated for 2 weeks with MXC-017 (5 i.p. injections per week, 50 mg/kg) or irradiated and treated with MXC-017. A dose of 4 Gy was chosen to ensure enough surviving cells at the end of the treatment course. Control animals were sham-irradiated and injected with solvent **(Fig. 3A)**. At the end of the two-week treatment period, tumors were explanted, digested into single cells and subjected to ELDAs. Both MXC-017 treatment alone and in combination with radiation reduced the number of spheres formed **(Fig. 3B/C)** and significantly decreased the *in vivo* GSC frequency **(Fig. 3D; Suppl. Tables 5 and 6)**. Furthermore, when HK-374-bearing NSG mice were treated with a single dose of 10 Gy to the tumor followed by continuous administration of MXC-017 (50 mg/kg), median survival increased significantly from 34 to 60 days (*p*<0.0001, Log-rank test; **Fig. 3E**) with no observed clinical toxicity and consistent weight gain (**Fig. 3F**). Note that the single fraction of 10 Gy is equivalent to a total dose of 18 Gy in 2 Gy fractions (for an alpha/beta ratio of 9 for GBM) and does not improve median survival in this PDOX model by itself. Immunostaining of the brain sections with an antibody against human mitochondria (**Fig. 3G**) or the neural stem cell marker nestin (**Fig. 3H**) indicated effective elimination of implanted human tumor cells after combined treatment. Furthermore, long-term surviving animals after combined treatment were tumor-free at the time of euthanasia (**Fig. 3I**).

**Figure 3.**
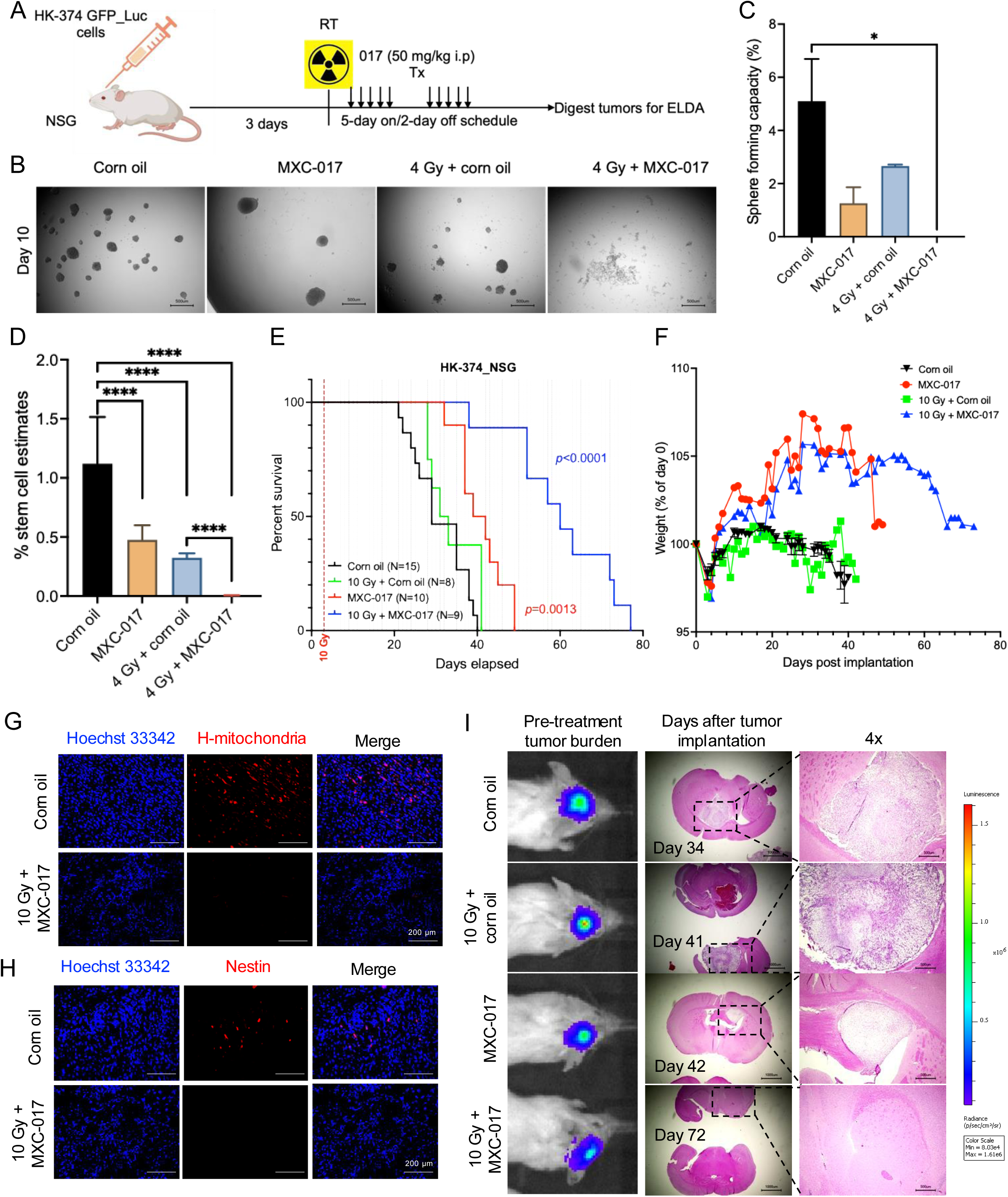
MXC-017 suppresses tumor self-renewal and prolongs survival in a PDOX GBM mouse model. **(A)** Experimental schematic illustrating the *in vivo* evaluation of MXC-017’s effects on GBM tumor self-renewal. **(B)** Representative images of tumor-derived spheres isolated from mouse brains after 2 weeks of treatment. **(C)** Quantification of sphere-forming capacity from tumor-derived cells. **(D)** Stem cell frequency analysis measured using ELDA software. **(E)** Kaplan–Meier survival curves of NSG mice intracranially implanted with HK-374-GFP-Luc patient-derived GBM cells. Tumors were allowed to graft for 3 days, followed by a single dose of 0 or 10 Gy radiation and treatment with either corn oil or MXC-017 (50 mg/kg, i.p., 5 days on/2 days off) until study endpoint. Dotted vertical lines indicate days of MXC-017 treatment. Survival comparisons were analyzed using the Log-rank (Mantel-Cox) test. The table below shows the number of mice alive at each time point. **(F)** Body weight curves of NSG mice in each treatment group throughout the study. **(G/H)** Immunofluorescence staining for human mitochondria and nestin in brain tissues from the 10 Gy + MXC-017 group versus corn oil controls. **(I)** Bioluminescence images showing initial tumor burden in long-term survivors, along with H&E staining of brain sections at euthanasia. Right panels display 4× magnification of the presumed tumor region. All experiments have been performed with at least 3 biological independent repeats. *P*-values were calculated using one-way ANOVA. * *p*-value < 0.05, **** *p*-value < 0.0001.

### MXC-017 binds to vimentin and prevents decompaction of vimentin intermediate filaments

Our HTS was an unbiased phenotypic screen and as such it did not identify hits against a specific molecular target. To directly determine the molecular target of MXC-017, we synthesized an alkyne-modified analog of MXC-017 (**Fig. 4A**), that could be bound to azide-tagged agarose beads via a click-chemistry reaction. Control experiments demonstrated that the alkyne-modified MXC-017 retained its biological activity, as evidenced by its ability to reduce sphere-forming capacity in GBM and decrease GSC frequencies (**Fig. 4B**).

**Figure 4.**
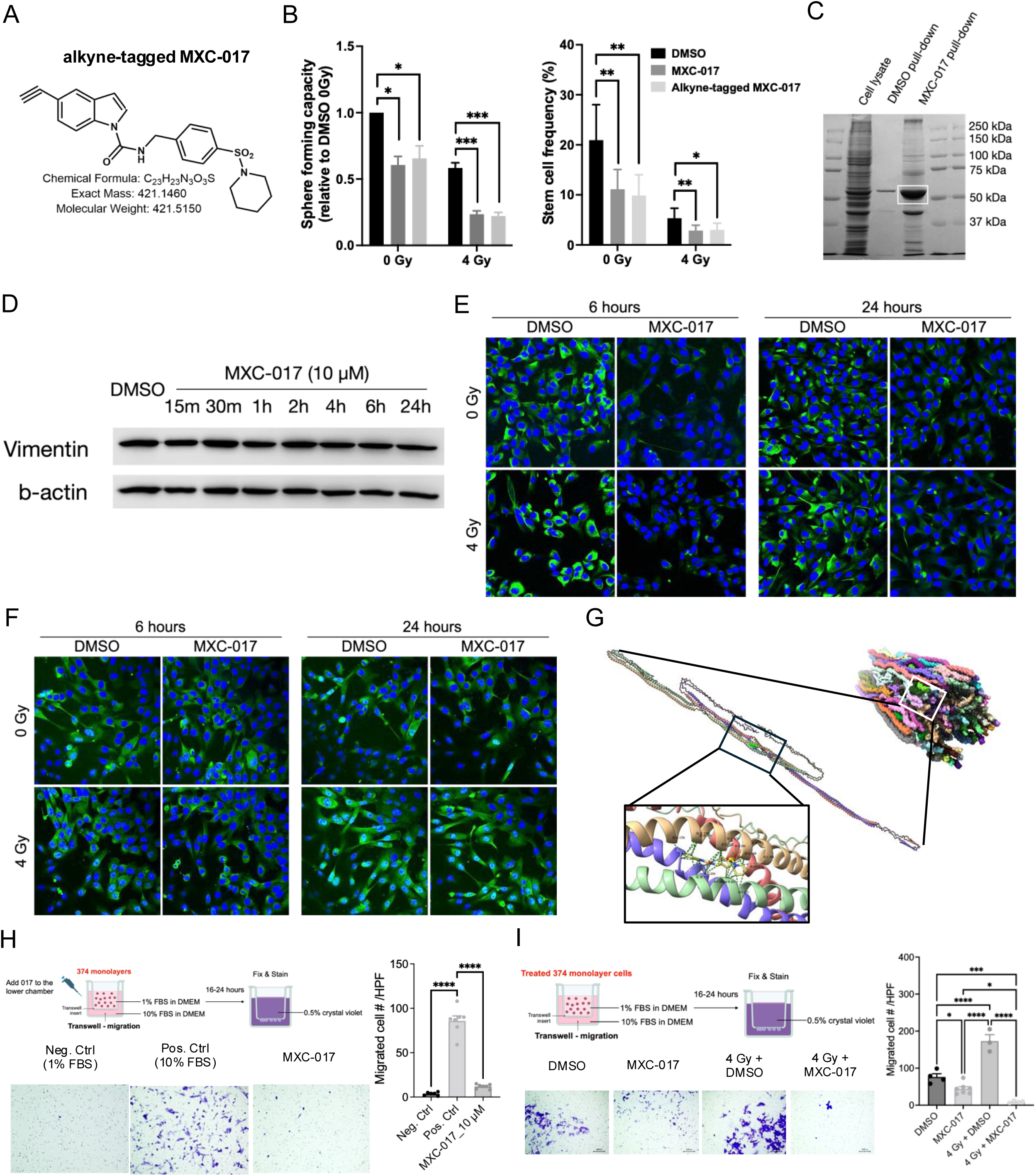
MXC-017 binds to vimentin and inhibits decompaction of vimentin intermediate filaments. **(A)** Chemical structure of alkyne-modified MXC-017. **(B)** Sphere-forming capacity and stem cell frequency of HK-374 cells treated with either MXC-017 or alkyne-modified MXC-017, with or without a single dose of 4 Gy radiation. **(C)** Coomassie-stained SDS-PAGE gel showing protein profiles from whole cell lysates, DMSO control pull-down, and alkyne-modified MXC-017 pull-down. The prominent band specific to MXC-017 pull-down is highlighted with a white box. **(D)** Western blotting of vimentin in HK-374 cells treated with 10 µM MXC-017 at various time points. **(E)** Confocal microscopy using a monoclonal antibody targeting the vimentin head domain (surrounding Arg45) in HK-374 cells treated with 10 µM MXC-017, with or without a single dose of 4 Gy radiation, imaged at 6 and 24 hours post-radiation. **(F)** Confocal microscopy using a polyclonal vimentin antibody under the same treatment condition as in **(E). (G)** *In silico* docking using the AutoDock Vina algorithm identified a potential MXC-017 binding site with vimentin pocket #45 with contact to the alpha-helices of all 4 vimentin molecules and a binding energy of -8.77 kcal/mol. **(H)** Transwell invasion assay of HK-374 cells with MXC-017 (10 µM) added to the lower chamber of an 8-µm pore insert. Media containing 1% FBS served as a negative control and 10% FBS as a positive control. Migrated cells were quantified per high-power field using Image J. **(I)** Invasion assay of HK-374 cell pretreated with MXC-017 (10 µM) for 48 hours with or without a single dose of 4 Gy radiation. Quantification of migrated cells per high-power field was performed using Image J. All experiments have been performed with at least 3 biological independent repeats. *P*-values were calculated using one-way ANOVA. * *p*-value < 0.05, ** *p*-value < 0.01, *** *p*-value < 0.001, **** *p*-value < 0.0001.

Gel-electrophoresis of proteins bound to MXC-017-tagged beads resulted in a prominent band (**Fig. 4C**) and subsequent mass spectroscopy identified vimentin as target for MXC-017 **(Suppl. Fig. 2A)**. Western blotting of protein from cells treated with MXC-017 using a polyclonal antibody showed no change in total vimentin levels over time **(Fig. 4D)** or changes in phosphorylation of Ser39, Ser56, or Ser83 **(Suppl. Fig. 2B)**. Confocal imaging of cells irradiated and/or treated with MXC-017 using a monoclonal antibody against a peptide in the head region of vimentin protein surrounding Arg45 showed the well-known induction of vimentin expression in response to radiation (30) **(Fig. 4E)**. Treatment with MXC-017 reduced detectable baseline levels of vimentin and prevented the radiation induced increase of the vimentin signal **(Fig. 4E)**, while confocal imaging using a polyclonal antibody against vimentin was consistent with the Western blot data, showing no change in total vimentin levels **(Fig. 4F)**.

Next, we performed *in silico* docking experiments using the MolModa software package vers. 1.0.1 (18). First, we used a truncated tetramer model from the PDB data base (PDB ID: 3KLT) previously used to study binding of the vimentin inhibitor ALD-R491, which reported a Kd of 328 ± 12.66 nM (31). *In silico* docking of MXC-017 using the AutoDock Vina algorithm (20) indicated a binding energy of -9.63 kcal/mol in the region where ALD-R491 binds to vimentin (**Suppl. Fig. 2C**), suggesting a Kd of 85.8nM for MXC-017 and thus, binding of MXC-017 to vimentin comparable to ALD-R491.

To validate these findings we obtained a recently published, more complete atomic model of vimentin from the PDB data base (8RVE) (19) to identify binding pockets in FPocketWeb (32). For the single tetramer comprising of a 2A-2B and a 1A-1B dimer derived from this model (19) FPocketWeb identified a total of 45 potential pockets with varying levels of druggability scores (**Suppl. Fig. 2D**). *In silico* docking using the AutoDock Vina algorithm (20) indicated a potential binding site for MXC-017 in pocket #45 with contact to the alpha-helices of all 4 vimentin molecules and a binding energy of -8.77 kcal/mol (**Fig. 4G; Suppl. Movie 1**).

Given the well-established role of vimentin in cytoskeletal remodeling and tumor cell invasion/migration, we next assessed the effect of MXC-017 on GBM cell migration using a transwell assay with two experimental approaches. In the first setup, MXC-017 was added to the lower chamber of the transwell system to assess its effect as a chemoattractant inhibitor. Here, we demonstrated that MXC-017 significantly reduced the migratory capacity of GBM cells, as evidenced by fewer cells penetrating through the membrane **(Fig. 4H)**. In the second approach, GBM cells were pre-treated with MXC-017, with or without a single dose of 4 Gy radiation, prior to seeding in the transwell assay. Pre-treated cells displayed markedly impaired migration compared to untreated or radiation-only controls **(Fig. 4I)**, indicating that MXC-017 suppresses GBM cell motility independently and in combination with radiation.

### MXC-017 has no off-target effects in GBM cells

To test for potential off target effects for MXC-017 we first performed metabolomics of HK-374 cells treated with radiation and/or MXC-017. The observed changes 48 hours after drug treatment were rather small and non-significant, thus excluding metabolic modulation as an additional mode of action **(Suppl. Fig. 3)**.

A whole kinome screen (Eurofins, France) reported less than 50% inhibition of Flt3, TNIK, and JNK3 by MXC-017 (10 µM) **(Suppl. Fig. 4A)** but in confirmatory experiments (Reaction Biology, PA) we could not establish an IC_50_ for MXC-017 for either of the kinases **(Suppl. Fig. 4B)**, thus suggesting that MXC-017 did not primarily act as a small molecule kinase inhibitor.

Next, we performed bulk RNA sequencing at 48h after drug treatment to detect global transcriptomic changes in response to MXC-017 treatment. The combination of radiation with MXC-017 compared to radiation alone only lead to 239 up- and 118 down-regulated differentially expressed genes (**Fig. 4A/B**) and no characteristic gene expression pattern, different from that of cells receiving only radiation (**Fig. 4C**). Gene set enrichment analysis revealed the typical hallmark gene sets upregulated after irradiation including the p53 pathway and pro-inflammatory responses. Compared to radiation alone, the combination of radiation and MXC-017 enhanced upregulation of KRAS signaling and the proinflammatory response while further downregulating E2F target genes and the G2M checkpoint (**Fig. 4D**). To assess whether the combination treatment affects cellular stemness, we analyzed the downregulated DEGs (any fold change with padj <0.05) in the combination group versus radiation alone using the StemChecker tool. Interestingly, we observed significant overlap between these downregulated genes and gene signatures associated with various stem cell types, including embryonic stem cells (272/2909 genes, padj = 9^-16^), embryonal carcinoma (63/628 genes, padj = 2.2^-11^), and hematopoietic (62/949 genes, padj = 4.1^-5^) stem cell types (**Fig . 4E/F**).

To assess if MXC-017 treatment changes the cellular composition of GBM we next performed scRNAseq in 4 PDOX lines. Gliomaspheres were plated onto poly-D-lysine/laminin-coated plates to prevent differentiating cells from dying from anoikis and harvested 5 days after irradiation. Using a gene program-centric meta-atlas of published transcriptomic studies (24) we annotated cell populations in HK-374 and HK-390 cells (classical TCGA subtype) and HK-217 and HK-244 cells (proneural TCGA subtype) and determined cell type proportions at baseline and in response to treatments with radiation and/or MXC-017. While cell type distributions differed between the different PDX lines at baseline, changes in response to MXC-017 alone or in combination with radiation were small (**Fig. 5G; Suppl. Fig. 5A-D; Suppl. Tables 7-11**), thus excluding elimination of specific cell types as an explanation for the effects of MXC-017. Likewise, when we applied a universal cellular senescence signature for scoring (25), there was no significant indication that MXC-017’s impact on GSCs was mediated through the induction of senescence (**Fig.5H**). Finally, we tested if MXC-017 would change the cell composition of tumors *in vivo*. HK-374 cells were injected into the brains of NSG mice. Tumors were treated, the cells harvested and analyzed with scRNAseq. While the overall cell composition changed when cells were grown *in vivo*, the addition of MXC-017 did not change the proportions of the cell types (**Suppl. Fig. 5E-J**).

**Figure 5.**
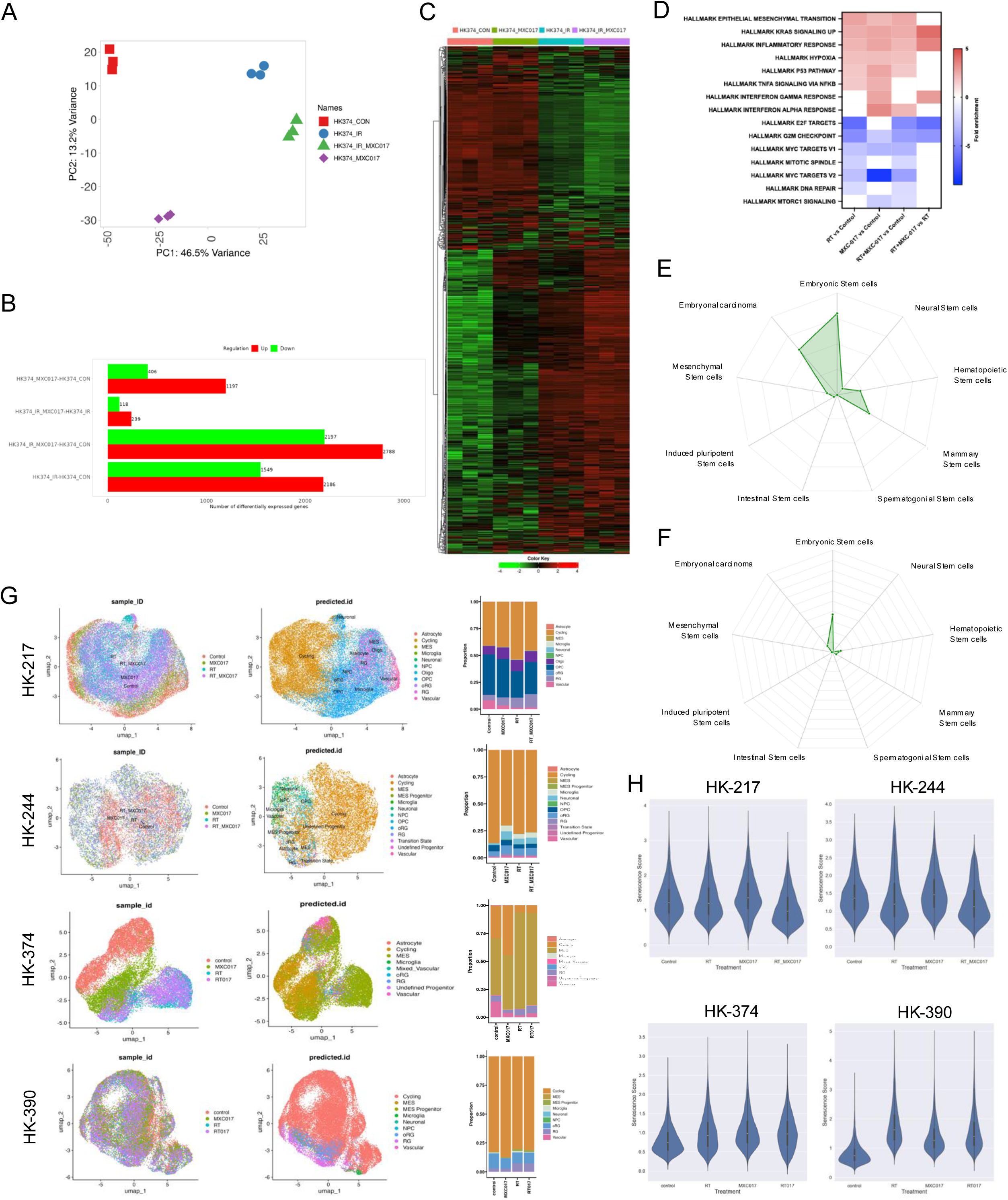
MXC-017 has no off-target effects in GBM cells. **(A)** Principal component analysis (PCA) of bulk RNA-sequencing data. **(B)** Bar graph displaying the number of differentially upregulated and downregulated genes identified across various group comparisons. **(C)** Heatmap with hierarchical clustering showing global gene expression profiles among treatment conditions. **(D)** Heatmap of significantly upregulated and downregulated signaling pathways based on gene set enrichment analysis. **(E/F)** Overlap analysis of downregulated DEGs in MXC-017/RT versus radiation alone comparison (any fold change, padj <0.05), revealing enrichment within gene expression signatures associated with various stem cell types, represented as -log10 p-value. **(G)** UMAP plots of clusters identified 5 days after initial of MXC-017 treatment across four experimental conditions, along with projected cell types annotations from all 4 PDOX lines. Stacked column graphs show the distribution of identified cell types by treatment group. (H) Violin plots illustrating senescence scores across treatment groups in all 4 PDOX lines.

### MXC-017 has low normal tissue toxicity in vitro and in vivo

The high efficacy of MXC-017 targeting GSCs and preventing phenotype conversion led us to investigate its potential cytotoxicity in non-cancerous cells. When we performed clonogenic survival assays using NIH3T3 fibroblasts and EOC20 microglia cells, we detected no significant toxicity (**Fig. 6A/B**). In contrast, normal human astrocytes showed a modest but statistically significant decrease in plating efficacy in the presence of MXC-017 (**Fig. 6C**). Importantly, primary murine neural stem/progenitor cells showed no evidence of toxicity when treated with MXC-017 alone or in combination with radiation, as assessed by sphere-forming assay (**Fig. 6D**).

**Figure 6.**
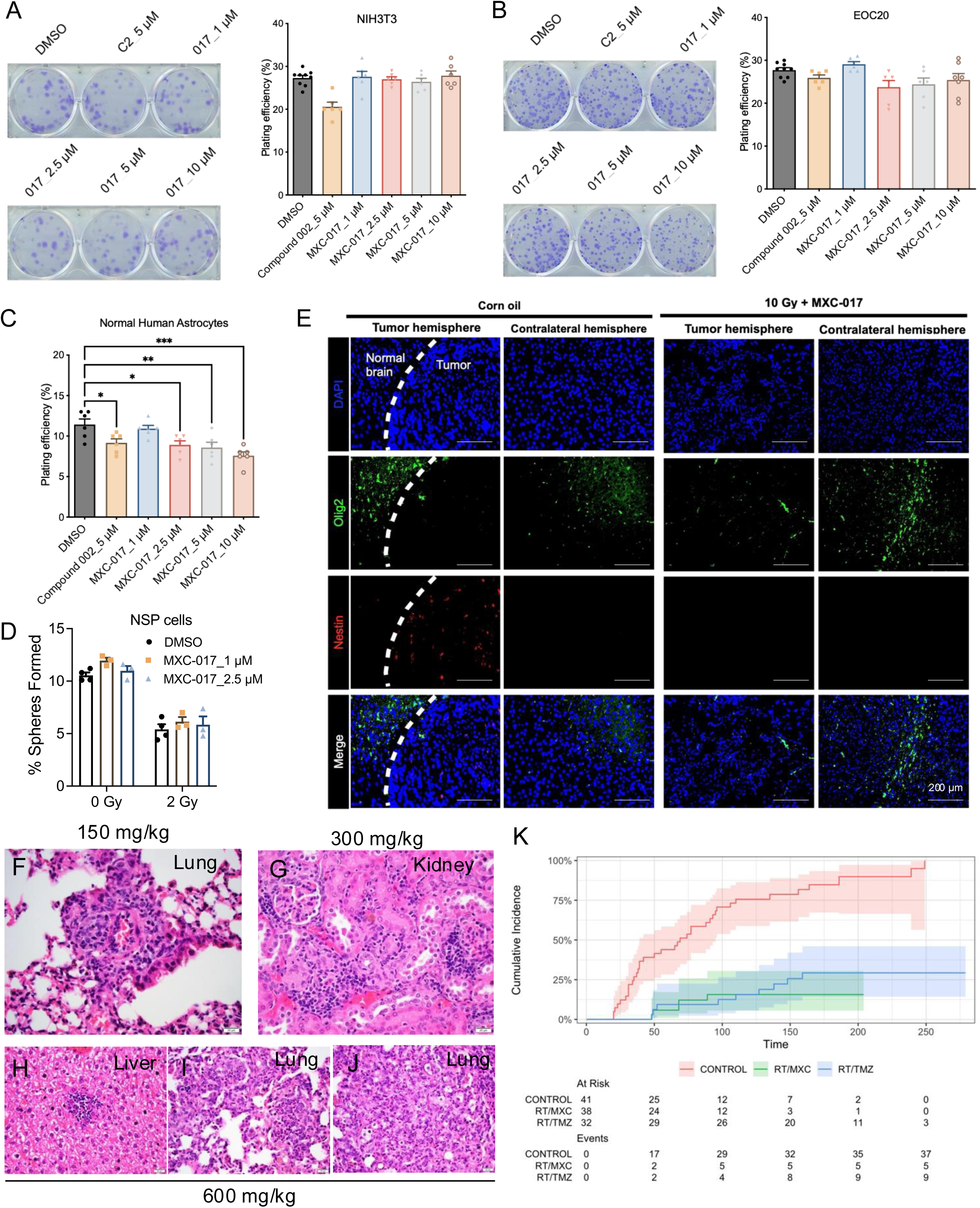
MXC-017 exhibits minimal toxicity to normal tissues in both *in vitro* and *in vivo* models. Representative images and quantification of colony formation in NIH3T3 fibroblasts following treatment with Compound 002 or MXC-017, assessed via clonogenic assay. **(A)** Clonogenic assay images and plating efficiency analysis of EOC20 microglial cells treated with Compound 002 or MXC-017. **(C)** Quantification of plating efficiency in normal human astrocytes treated with Compound 002 or MXC-017, using a clonogenic assay. **(D)** Sphere-forming capacity of primary murine neural stem/progenitor cells treated with MXC-017 alone or in combination with a single 2 Gy dose of radiation. **(E)** Immunofluorescence imaging of Olig2 and Nestin expression in both the tumor-bearing and contralateral hemispheres of HK-374 tumor–bearing NSG mice treated with either 10 Gy plus MXC-017 or corn oil vehicle control. **(F)** H&E-stained lung section from C57BL/6 mice treated with MXC-017 at 150 mg/kg, showing mild perivascular inflammation and superficial vasculitis. **(G)** H&E image of kidney tissue from mice treated with 300 mg/kg MXC-017, indicating focal interstitial inflammation in the renal cortex. **(H)** Liver section from mice treated with 600 mg/kg MXC-017, displaying focal lobular inflammation and evidence of hepatocellular necrosis. **(I/J)** H&E-stained lung sections from mice treated with 600 mg/kg MXC-017, revealing neutrophilic infiltration in the alveolar spaces and areas of dense inflammation. **(K)** Cumulative incidence plot of tumor-related mortality across 17 different patient-derived orthotopic xenograft (PDOX) GBM models. Survival outcomes for MXC-017 combined with radiation were benchmarked against a standard treatment regimen of one cycle of temozolomide (TMZ) plus radiation, and both were compared to untreated control groups. All experiments have been performed with at least 3 biological independent repeats. *P*-values were calculated using one-way ANOVA. * *p*-value < 0.05, ** *p*-value < 0.01, *** *p*-value < 0.001.

To further investigate potential CNS toxicity *in vivo*, we treated HK-374 tumor-bearing NSG mice with MXC-017 (50 mg/kg i.p. 5 days on – 2 days off for two weeks) with a single dose of 10 Gy radiation at day 3 post-implantation. Brains were harvested and immunostained for nestin and olig2 to assess effects on tumor cells and normal neural populations, respectively. While the combination of MXC-017 and radiation effectively eliminated nestin-positive human tumor cells, no loss of olig2-positive oligodendrocytes was observed in the contralateral (non-tumor-bearing) hemisphere, suggesting MXC-017’s selective anti-tumor activity with minimal damage to normal brain tissue (**Fig. 6E**).

Finally, we determined the maximum tolerated dose for MXC-017 in mice. Male and female C57Bl/6 mice were treated a single dose of MXC-017 at 150, 300, 600 and 900 mg/kg. Animals were observed for 2 weeks after which a necropsy was performed, major organs and were harvested. None of the animals showed signs of toxicity. A second cohort of males and female mice was treated with 5 weekly doses of MXC-017 at 150, 300 and 600 mg/kg for 2 weeks. None of the animals show clinical signs of toxicity. Forty-eight hours after the last injection, major organs (heart, lung, liver, kidneys, spleen, and brain), whole blood and plasma were collected. None of the animals showed significant hematological or biochemical abnormalities (**Suppl. Figs. 6-8**). At 150 mg/kg, we observed mild lung toxicity characterized by perivascular inflammation and superficial vasculitis (**Fig. 6F**). At 300 mg/kg, focal interstitial inflammation was detected in the kidney cortex (**Fig. 6G**). At 600 mg/kg, the liver showed focal lobular inflammation and hepatocellular necrosis (**Fig. 6H**). In the lungs, there were signs of acute pneumonia, with predominantly neutrophilic inflammation in the alveolar spaces (**Fig. 6I**), along with areas of dense inflammation featuring admixed neutrophils and a prominent histiocytic, vaguely granulomatous reaction (**Fig. 6J**).

Based on this dose escalation study we determined the MTD at 150 mg/kg. Using 17 different PDOXs in male and female NSG mice, covering the proneural, classical and mesenchymal TCGA subtype we performed a survival experiment benchmarking MXC-017 in combination with radiation against one cycle of TMZ (33) plus irradiation and compared both to untreated control animals. Because MXC-017 was given continuously for more than 140 days, thus increasing the probability for opportunist infections in highly immunodeficient NSG mice, we used a competing risk model (34) to calculate the cumulative incidences for tumor-related death (**Fig. 6K**). MXC-017 in combination with radiation significantly reduced the probability for tumor related death compared to control animals. MXC-017 showed a trend for reduced probability for tumor related death compared to TMZ combined with radiation, but the difference did not reach the level of statistical significance. Yet, while TMZ is known for its significant side effects (35), MXC-017 was well tolerated, even when given until the animals reached euthanasia endpoints.

## Discussion

GBM continues to be one of the most difficult to treat tumors with low median survival und unacceptable low long-term survival. Radiotherapy against GBM has been iterated though all available technologies and fractionation schemes (36) and while it remains the most powerful treatment modality next to surgery, its contribution to treatment outcome has stagnated. Likewise, chemotherapy, targeted therapies, and biologics have not improved outcome (37) and surgery, radiotherapy, and temozolomide remain the standard-of-care.

Previously, we reported radiation-induced cellular plasticity and the generation of GSCs from non-stem glioma cells as drivers of treatment failure in GBM (9,10,12,38–40). Here, we used results from our unbiased, phenotypic HTS (26) of libraries containing over 83,000 compounds to identify candidates that target existing GSCs, prevent the radiation-induced conversion of non-stem GBM cells into induced GSCs, while not working as classical radiosensitizers. We limited the resulting hits to compounds with no Lipinsiky violations (27) and a CNS Multiparameter Optimization Score (41) of >=4, to predict BBB penetration, a major hurdle in drug development against GBM. Based on our hits we synthesized 114 novel, structure-related analogs that target existing GSCs and their *de novo* generation. Our lead compound MXC-017 showed a level of BBB-penetration in mice that exceeded that of the psychotropic anti-schizophrenia drug quetiapine in our model system (9).

MXC-017 significantly reduced self-renewal capacity and GSC frequencies in a panel of IDHwt GBM lines of classical, proneural and mesenchymal TCGA subtypes *in vitro* and doubled the median survival in PDOX models *in vivo*. While it showed a trend of outperforming TMZ in combination with radiation it was most notable that MXC-017 caused very little if any normal tissue toxicity at the determined MTD of 150mg/kg. Because molecular its target, vimentin, is overexpressed in GBM (42), MXC-017 appears to offer a wide therapeutic window, even when combined with radiation.

Our observation that binding of an antibody against a residue in the head region of the vimentin protein, folded into the inner lumen of the assembled filament, was reduced by MXC-017, suggests that MXC-017 locks vimentin in an assembled state, thereby impairing cell cycle progression, migration, stemness, phenotype plasticity and inducing apoptosis. A previous study reported radiation-induced expression of vimentin and increased vimentin-dependent migration in GBM (43). Here, we confirmed this effect of radiation on vimentin expression and migration demonstrate that MXC-017 effectively reversed this effect of radiation. This mode of action of MXC-017 on vimentin differs from other known vimentin-interacting compounds, which either down-regulate its expression (44,45) or cause its disassembly (45–48), yet all show anti-tumor activity, thereby establishing vimentin as a target for cancer therapy. Most of the compounds that affect vimentin expression or stability (45) have a wide range of other effects and vimentin is not the main target, thus limiting their application against tumors and for the two more specific vimentin-interacting compounds, FiVe1 (47) and ALD-R491 (48), it is unclear to which extent they can cross the BBB.

In GBM, high expression of vimentin is associated with progression and poor prognosis and targeting of vimentin using withaferin *in vitro* reduced migration of established GBM lines and diminished their viability (49). Furthermore, antibody targeting of cell surface vimentin (50) sensitized GSCs to TMZ and prolonged survival in orthotopic mouse models of GBM. However, in this study the GBM cells were preincubated with the antibody *ex vivo* (51), thus emphasizing the need for novel BBB-penetrating vimentin-interacting compounds.

Our HTS approach differed from most previous attempts to identify novel anti-GBM compounds in that it was an unbiased, phenotypic screen and the identification of vimentin as the molecular target was rather unexpected. With its lack of enzymatic activity, vimentin-targeting compounds are typically not intentionally discovered in HTSs.

Previous studies (51), reporting sensitization of GBM cells to TMZ by vimentin-targeting compounds and our data presented in the present study suggest that MXC-017 could enhance the efficacy of the current standard-of-care, radiotherapy plus TMZ. With its excellent antitumor effects in GBM *in vivo*, its ability to cross the BBB, lack of off-target effects or normal tissue toxicity, MXC-017 shows great translational potential for clinical use in combination with radiation in patients suffering from GBM.

## Author contributions

FP and MEJ conceived of the study. XC and HD synthesized the MXC compounds. LH performed most of the experiments. LA, AMC, MM, CC, ST, and CH performed some of the experiments. LLM, HIK, and RD provided materials. GE and JW performed mass spec for the PK data. ZZ and GL performed the competing risk model analysis. JEZ performed the histopathology. FP, LH, DA, and AB analyzed the data. LH and FP drafted the manuscript. All authors contributed to and approved the final version of the manuscript.

## Data and Material Availability

All data are included in the article and/or *SI Appendix.* All cell lines will be made available upon reasonable request to the corresponding author.

## Acknowledgements

This work used computational and storage services associated with the Hoffman2 Shared Cluster provided by UCLA Office of Advanced Research Computing’s Research Technology Group. This work was made possible, in part, through access to the following: The UCLA Broad Stem Cell Research Center Microscopy, Flow Cytometry, and Sequencing Cores, the UCLA Translational Pathology Core Laboratory, the UCLA Mass Spectrometry and Proteomics Laboratory, the UCLA Crump Preclinical Imaging Technology Center, the UCLA Metabolomics and Proteomics Shared Resource, the UCLA Molecular Screening Shared Resource, and the NIH Cancer Center Support Grant (2 P30 CA016042-44).

FP was supported by grants from the *National Cancer Institute* (R01CA260886, R01CA281682) and the UCLA Eli and Edythe Broad Center of Regenerative Medicine and Stem Cell Research Award Program. FP, MEJ and HIK were supported by the California Institute for Regenerative Medicine (CIRM; DISC2-14083). AB, HIK and FP were supported by the American Cancer Society (<u>CSCC-Team-23-980262-01-CSCC</u>). LLM, HIK, and FP were supported by the *National Cancer Institute* UCLA SPORE in Brain Cancer 5P50CA211015.

