## Supplemental Materials for "Design and Biological Activity of a Novel Brain Penetrant Urea Compound Against Glioblastoma"

Table of contents:

1. Supplementary Figures

2. Supplementary Tables

3. Supplementary Materials and Methods

4. References

#### Supplementary Figures

##### Supplementary Figure 1.

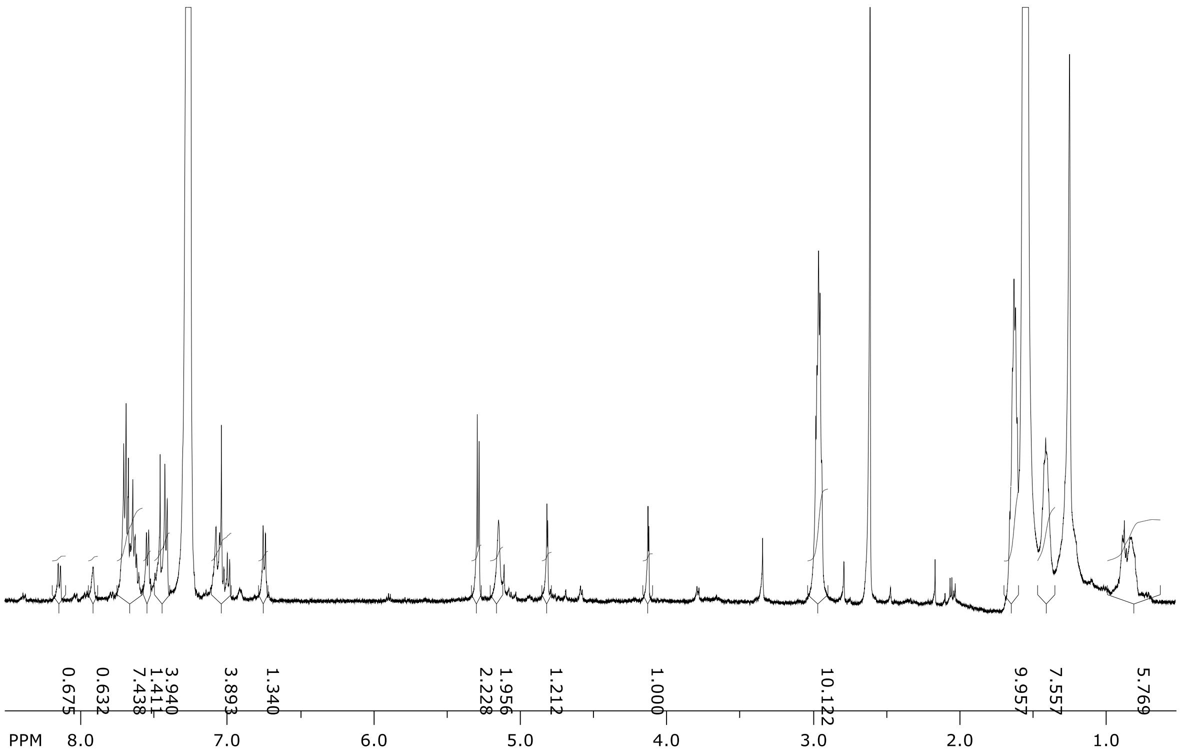


**Supplementary Figure 1. NMR spectroscopy analysis of Compound 002 (Molport-009-476-782).**

Proton nuclear magnetic resonance ($^1H$ NMR) spectroscopy of the original Compound 002 reveals multiple distinct peaks, indicating low compound purity and the presence of multiple degradation products.

**Supplementary Figure 2.**

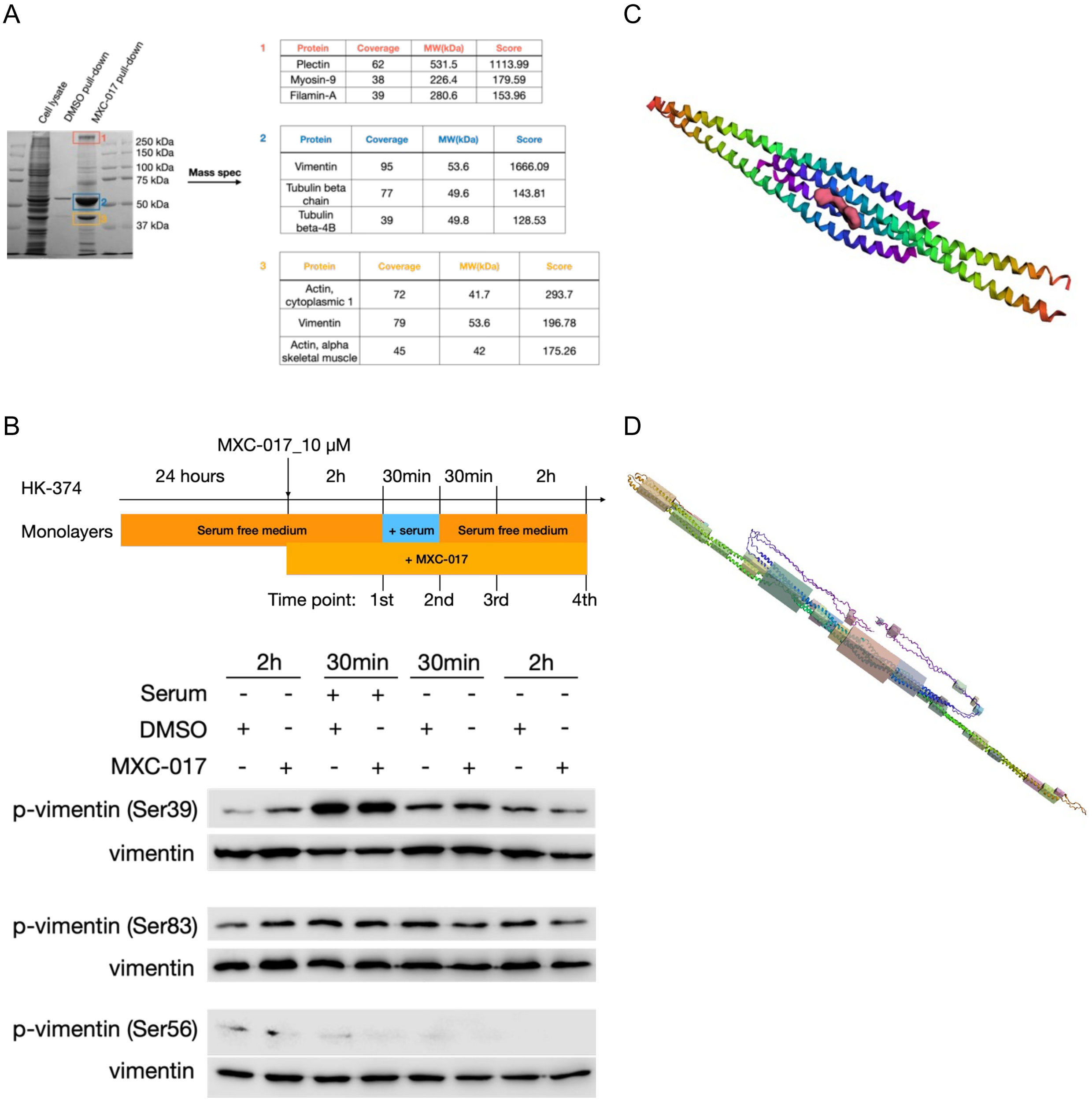


**Supplementary Figure 2. MXC-017 binds to vimentin and inhibits decompaction of vimentin intermediate filaments.**

**(A)** Coomassie-stained SDS-PAGE gel showing protein profiles from whole cell lysates, DMSO control pull-down, and alkyne-modified MXC-017 pull-down. Top 3 candidates from each prominent band from alkyne-modified MXC-017 pull-down gel using mass spectrometry. (B) Western blotting of p-vimentin at Ser 39, Ser 56, and Ser 83 sites and total vimentin. (C/D) *In silico* docking using the AutoDock Vina algorithm identified a potential MXC-017 binding site with vimentin pocket #45 with contact to the alpha-helices of all 4 vimentin molecules and a binding energy of -8.77 kcal/mol.

### Supplementary Figure 3.

### Metabolomics

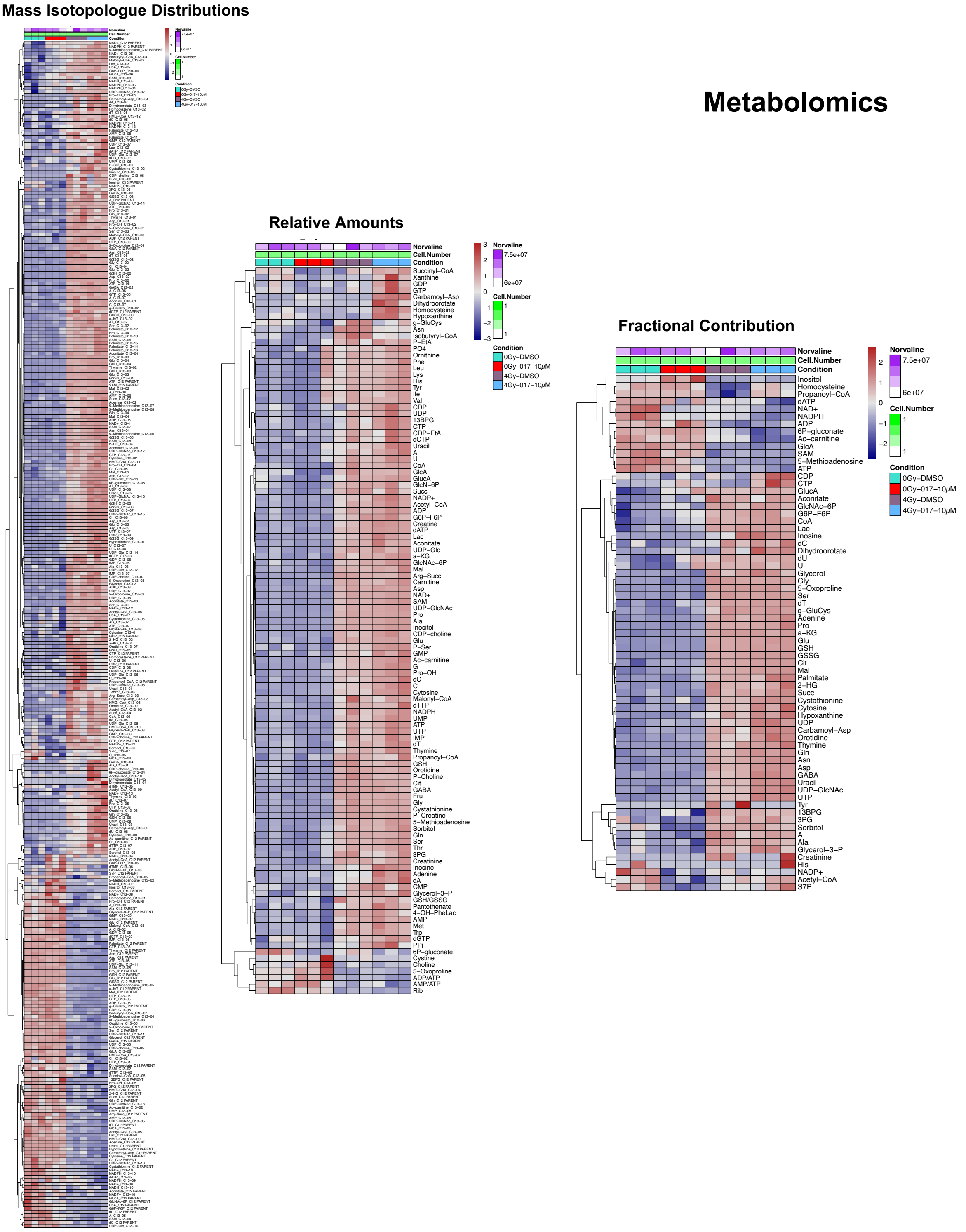

**Supplementary Figure 3. Metabolic profiling of HK-374 cells treated with MXC-017 (10 μM), with or without a single dose of 4 Gy radiation, assessed 24 hours post-treatment.**

### Supplementary Figure 4.

A

MCX017 Kinases inhibited

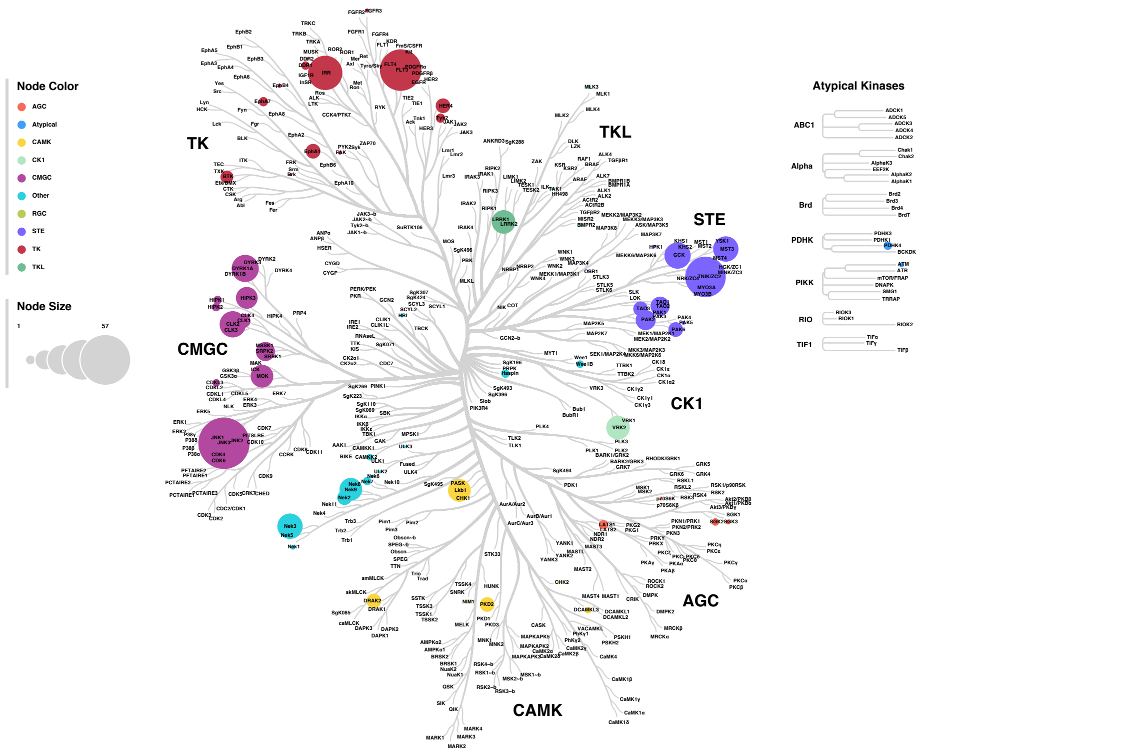


B

Compound IC50 Data for FLT3

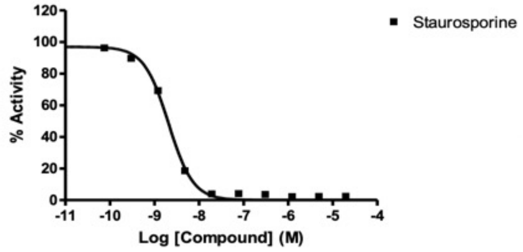


| | Staurosporine |
|---|---|
| HILLSLOPE | -1.582 |
| EC50 | 2.064e-009 |

Compound IC50 Data for JNK3

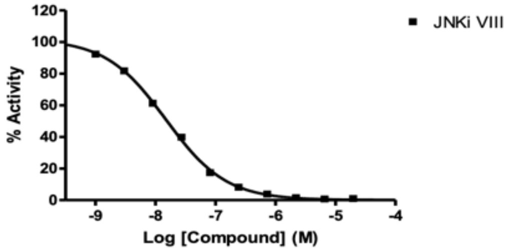


| | JNKi VIII |
|---|---|
| HILLSLOPE | -0.8682 |
| EC50 | 1.486e-008 |

Compound IC50 Data for TNIK

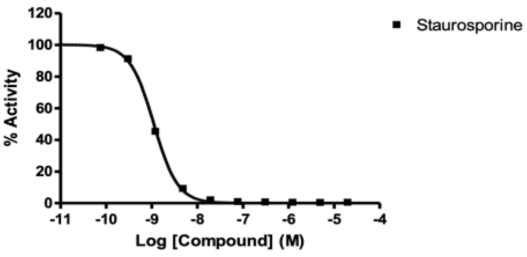


| | Staurosporine |
|---|---|
| HILLSLOPE | -1.664 |
| EC50 | 1.110e-009 |

| Kinase: | [ATP](μM): | Compound IC50* (M): | | IC50 (M) Control Cmpd | Control Cmpd ID |
|---|---|---|---|---|---|
| | | MXC017 | MXC017 | | |
| FLT3 | 50 | | | 2.06E-09 | STAUROSPORINE |
| JNK3 | 2.5 | not | detected | 1.49E-08 | JNKI VIII |
| TNIK | 50 | | | 1.11E-09 | STAUROSPORINE |

**Supplementary Figure 4. Whole kinome profiling (A) and a Kinase HotSpot assay (B) evaluating MXC-017 activity across a panel of kinase in HK-374 cells.**

**Supplementary Figure 5.**

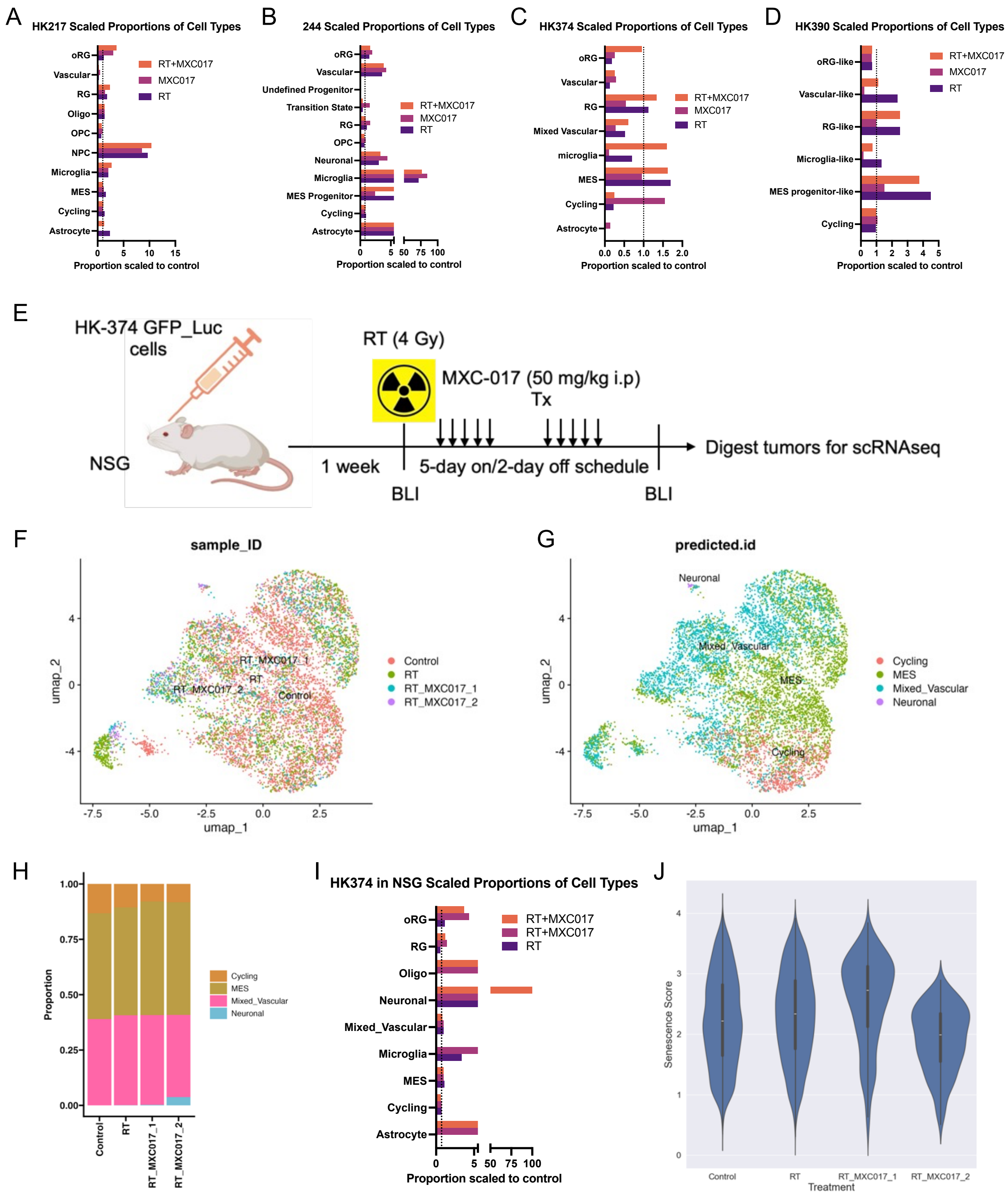

**Supplementary Figure 5. Single-cell RNA sequencing analysis.**

**(A-D)** Scaled cell type proportions relative to control across all treatment groups in in vitro single-cell RNA-seq datasets for HK-217, HK-244, HK-374, and HK-390. **(E)** Schematic overview of the experimental design for in vivo single-cell RNA-seq of HK-374. **(F/G)** UMAP plots showing identified clusters and projected cell type annotations from the HK-374 *in vivo* dataset. **(H/I)** Stacked column charts representing cell type composition and their relative proportions, scaled to control. **(J)** Violin plots illustrating senescence scores across treatment groups.

### Supplementary Figure 6.

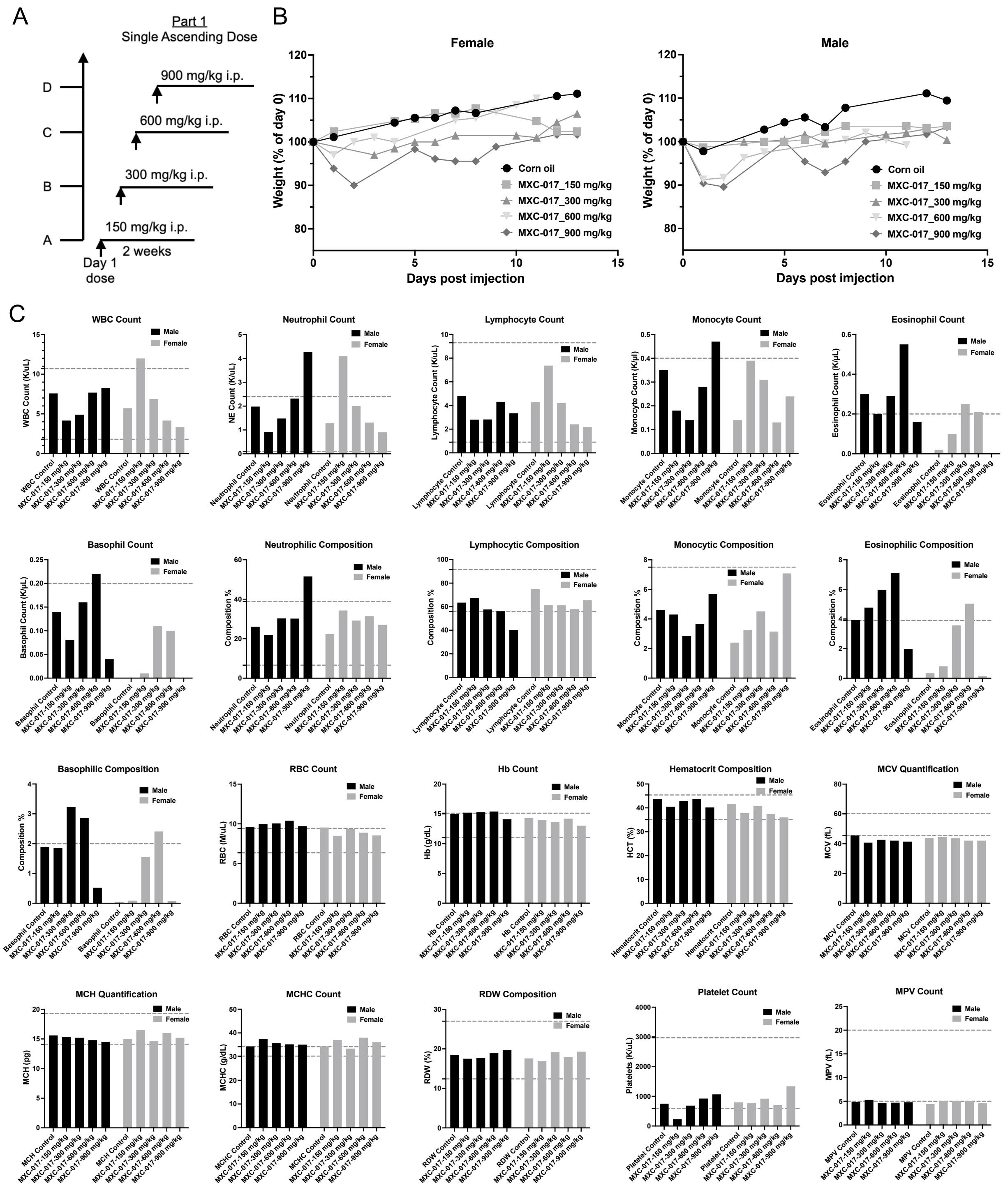

**Supplementary Figure 6. Hematological assessment from single-dose escalation MTD study using HemaVet® 950FS.**

**(A)** Dose escalation scheme for single-dose MXC-017 treatment. **(B)** Weight curves of treated animals across the dose escalation study. **(C)** Hematological parameters measured using the HemaVet®950FS analyzer.

### Supplementary Figure 7.

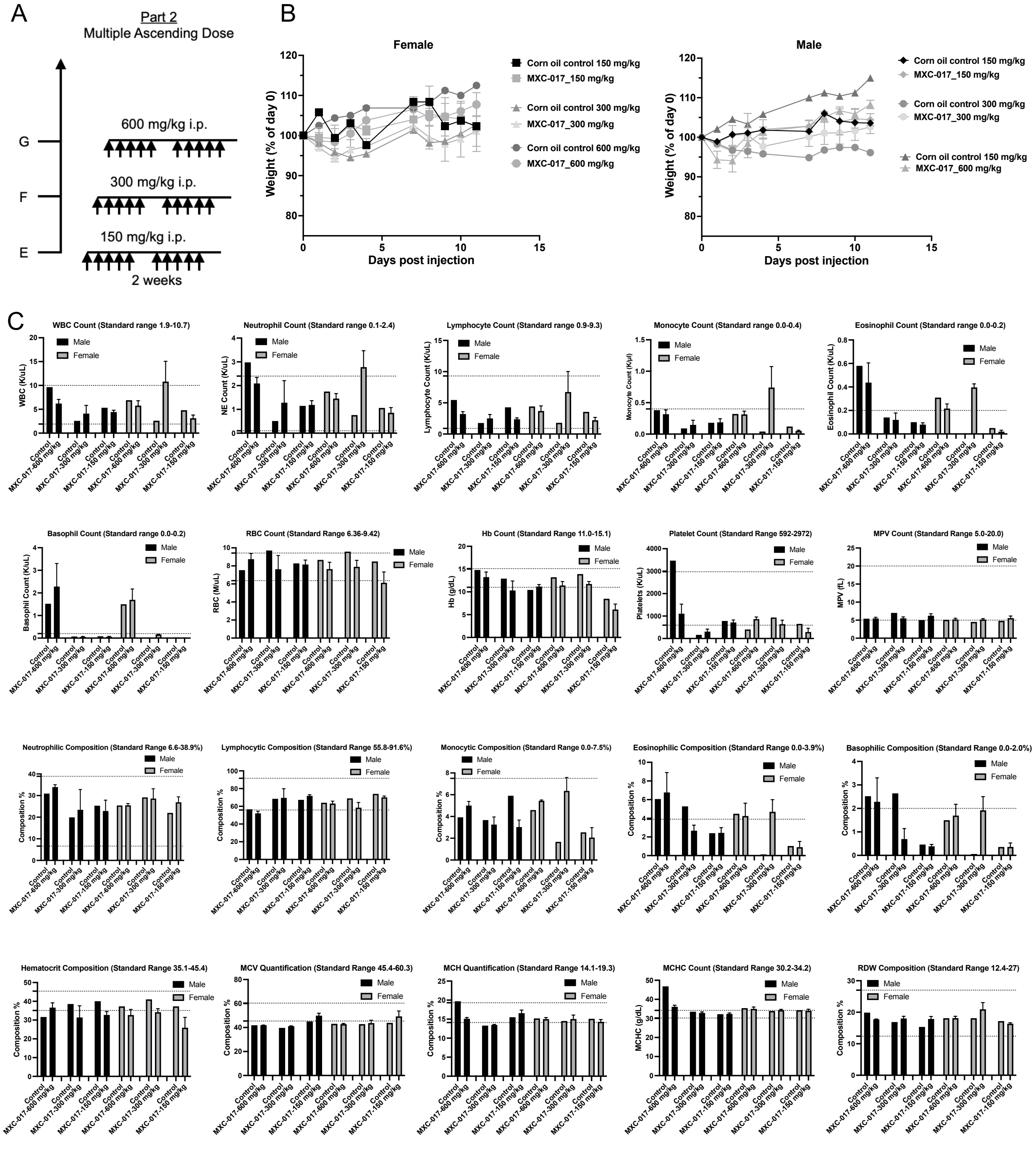

**Supplementary Figure 7. Hematological assessment from multipe-dose escalation MTD study using HemaVet® 950FS.**

**(A)** Dose escalation scheme for multiple-dose MXC-017 treatment. **(B)** Weight curves of treated animals across the dose escalation study. **(C)** Hematological parameters measured using the HemaVet®950FS analyzer.

#### Supplementary Figure 8.

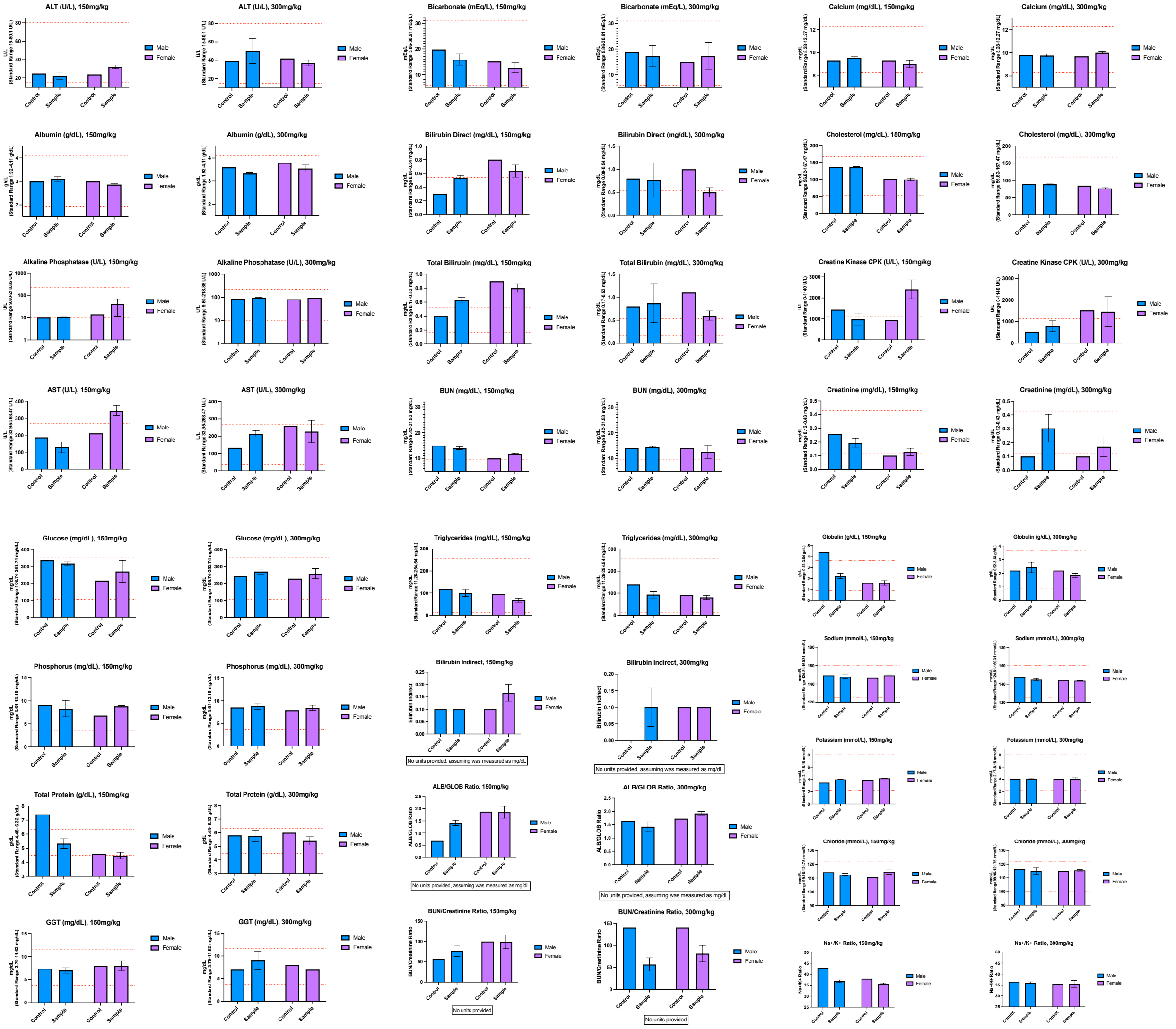

**Supplementary Figure 8. Comprehensive Plus Clinical Chemistry Panel analysis of serum samples from animals treated with multiple dose of MXC-017.**

#### Supplementary Tables

##### Supplementary Table 1. Patient demographics and TCGA-classification of GBM subtypes.

| Line | Origin | TCGA subtype | Culture P53 CN | EGFRvIII | PTEN | MGMT |
|---|---|---|---|---|---|---|
| HK-374 | Primary GBM | c | Loss "mosaic" | Positive | Positive | not methylated |
| HK-157 | Primary GBM | p | wt | Negative | Positive | Unknown |
| HK-308 | Recurrent GBM | m | Unknown | Positive | Positive | not methylated |
| HK-146 | Recurrent GBM | p | Unknown | Unknown | Unknown | Unknown |
| HK-385 | Unknown | p | Unknown | Unknown | Unknown | Unknown |
| HK-177 | Primary GBM | m | Unknown | Unknown | Unknown | Unknown |
| HK-378 | Recurrent GBM | c | Unknown | Negative | Unknown | not methylated |
| HK-248 | Recurrent GBM | m | CN loss | Negative | Positive | Unknown |
| HK-254 | Recurrent GBM | c | Unknown | Unknown | Unknown | Unknown |
| HK-390 | Primary GBM | c | wt | Negative | Positive | not methylated |
| HK-244 | Primary GBM | c | Unknown | Negative | Positive | not methylated |
| HK-336 | Recurrent GBM | m | CN loss | Negative | Positive | methylated |
| HK-339 | Recurrent GBM | m | Unknown | Unknown | Unknown | Unknown |
| HK-235 | PNET Grade IV | p | Unknown | Negative | Positive | Unknown |
| HK-217 | Primary GBM | p | wt | Negative | Negative | Unknown |
| HK-345 | Recurrent GBM | m | wt | Negative | Positive | not methylated |
| HK-393 | Recurrent GBM | m | Unknown | Unknown | Unknown | Unknown |

Abbreviations for TCGA subtypes: c: classical; m: mesenchymal; p: proneural.

**Supplementary Table 2. The chemical structures and molecular weights of lead compound 002 and its 114 analogs**

| Compound | Structure | MW |
|---|---|---|
| Compound 002 | 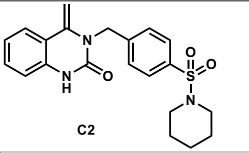 C2 | 397.49 |
| MXC-001 | 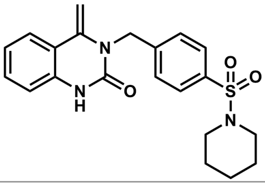 | 460.56 |
| MXC-002 | 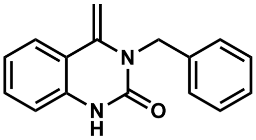 | 312.37 |
| MXC-003 | 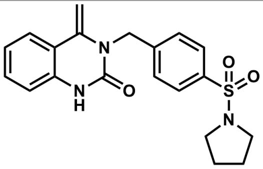 | 446.53 |
| MXC-004 | 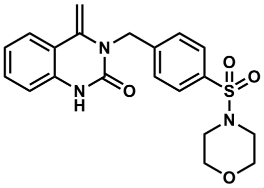 | 462.53 |
| MXC-005 | 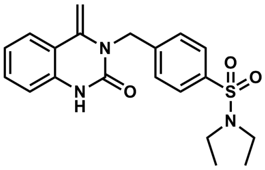 | 385.48 |
| MXC-006 | 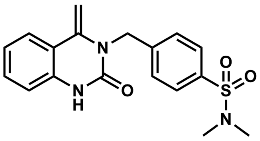 | 474.58 |
| MXC-007 | 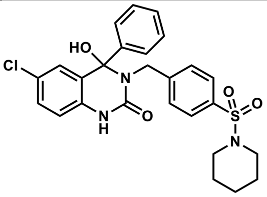 | 512.02 |
| MXC-008 | 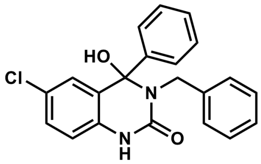 | 364.83 |

| | | |
|---|---|---|
| MXC-009A | 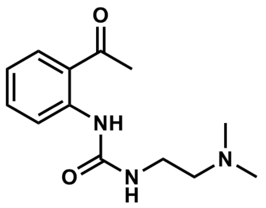 | 249.31 |
| MXC-009B | 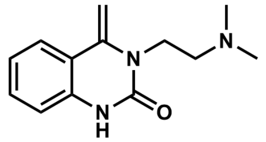 | 231.30 |
| MXC-010 | 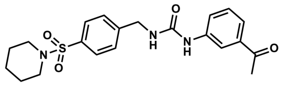 | 415.51 |
| MXC-011 | 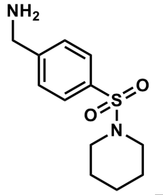 | 254.35 |
| MXC-012 | 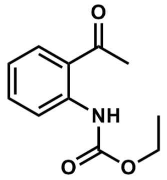 | 207.23 |
| MXC-013 | 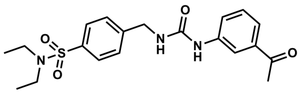 | 403.50 |
| MXC-014 | 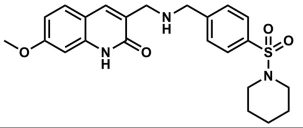 | 441.55 |
| MXC-015 | 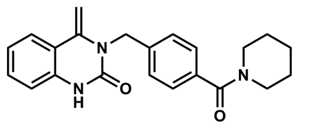 | 361.45 |
| MXC-016 | 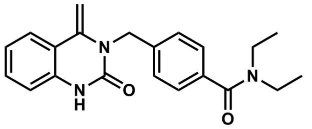 | 349.43 |
| MXC-017 | 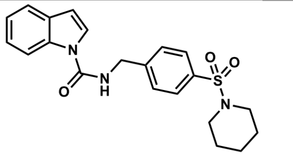 | 397.49 |
| MXC-018 | 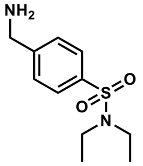 | 242.34 |

| | | |
|---|---|---|
| MXC-019 | 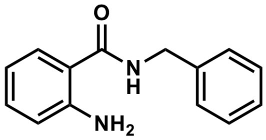 | 226.28 |
| MXC-020 | 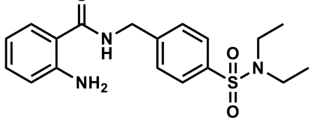 | 361.46 |
| MXC-021 | 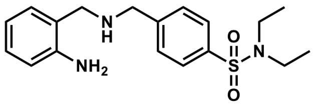 | 347.48 |
| MXC-022 | 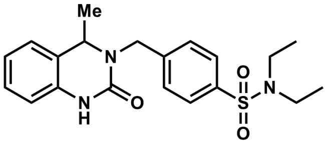 | 387.50 |
| MXC-023 | 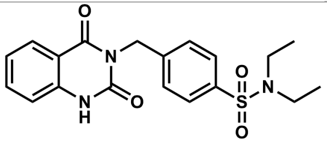 | 387.45 |
| MXC-024 | 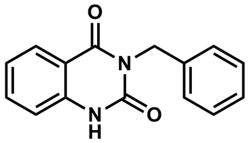 | 252.27 |
| MXC-025 | 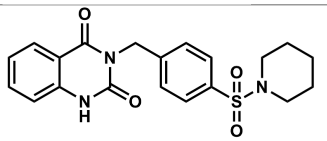 | 399.47 |
| MXC-026 | 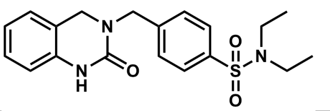 | 373.47 |
| MXC-027 | 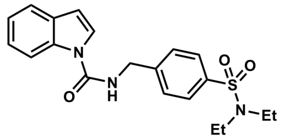 | 385.48 |
| MXC-028 | 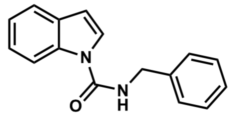 | 250.30 |
| MXC-029 | 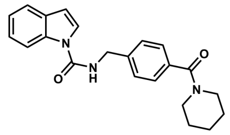 | 361.45 |
| MXC-030 | 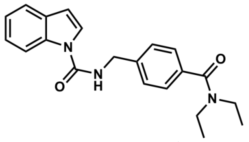 | 349.43 |

| | | |
|---|---|---|
| MXC-031 | 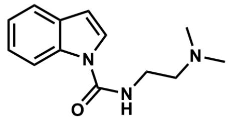 | 231.3 |
| MXC-032 | 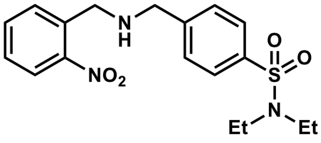 | 377.46 |
| MXC-033 | 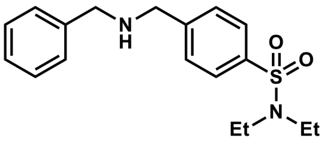 | 332.46 |
| MXC-034 | 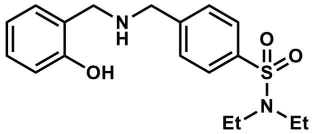 | 348.46 |
| MXC-035 | 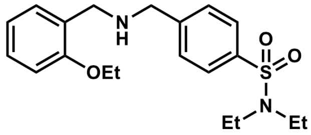 | 376.52 |
| MXC-036 | 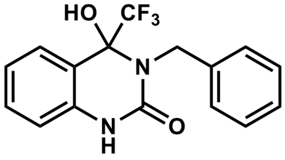 | 322.29 |
| MXC-037 | 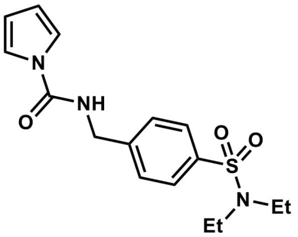 | 335.42 |
| MXC-038 | 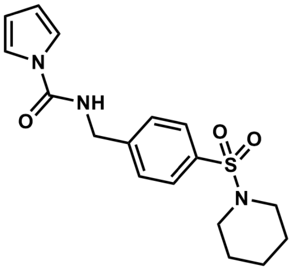 | 347.43 |
| MXC-039 | 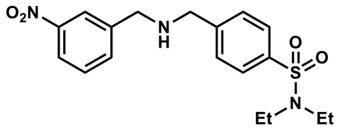 | 377.46 |

| | | |
|---|---|---|
| MXC-040 | 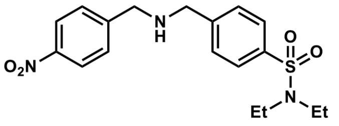 | 377.46 |
| MXC-041 | 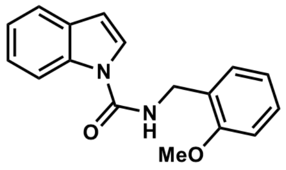 | 280.33 |
| MXC-042 | 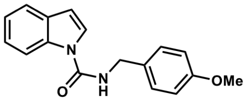 | 280.33 |
| MXC-043 | 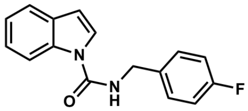 | 268.29 |
| MXC-044 | 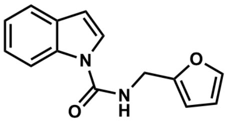 | 240.26 |
| MXC-045 | 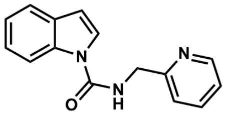 | 251.29 |
| MXC-046 | 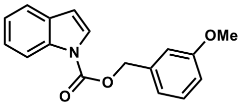 | 281.31 |
| MXC-047 | 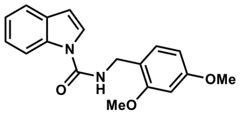 | 310.35 |
| MXC-048 | 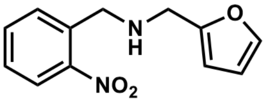 | 232.24 |
| MXC-049 | 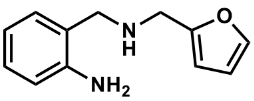 | 202.26 |
| MXC-050 | 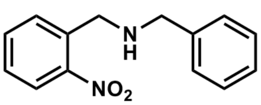 | 242.28 |
| MXC-051 | 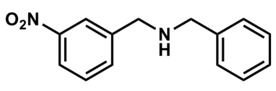 | 242.28 |
| MXC-052 | 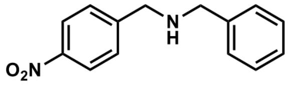 | 242.28 |
| MXC-053 | 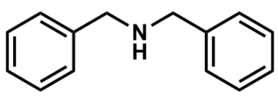 | 197.28 |

| | | |
|---|---|---|
| MXC-054 | 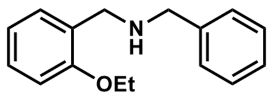 | 241.33 |
| MXC-055 | 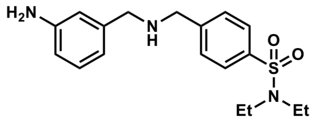 | 347.48 |
| MXC-056 | 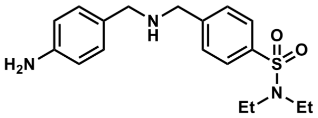 | 347.48 |
| MXC-057 | 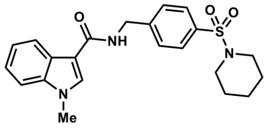 | 411.52 |
| MXC-058 | 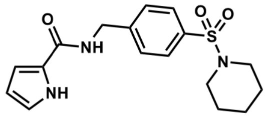 | 347.43 |
| MXC-059 | 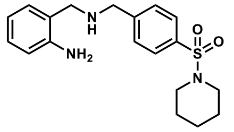 | 359.49 |
| MXC-060 | 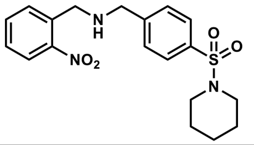 | 389.47 |
| MXC-061 | 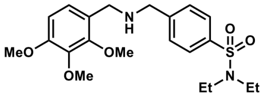 | 422.54 |
| MXC-062 | 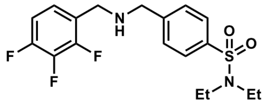 | 386.43 |
| MXC-063 | 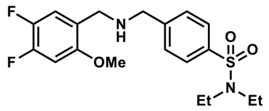 | 398.47 |
| MXC-064 | 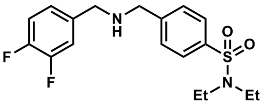 | 368.44 |
| MXC-065 | 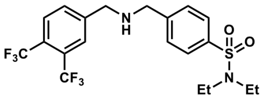 | 468.46 |
| MXC-066 | 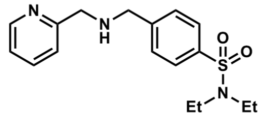 | 333.45 |

| | | |
|---|---|---|
| MXC-067 | 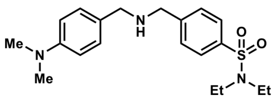 | 375.53 |
| MXC-068 | 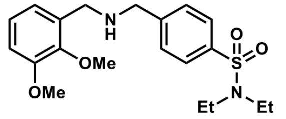 | 392.51 |
| MXC-069 | 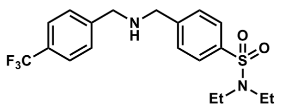 | 400.46 |
| MXC-070 | 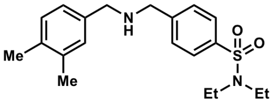 | 360.52 |
| MXC-071 | 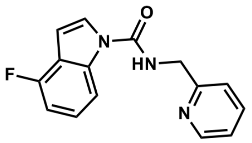 | 269.28 |
| MXC-072 | 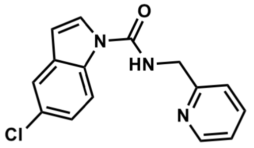 | 285.73 |
| MXC-073 | 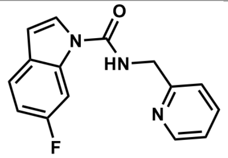 | 269.28 |
| MXC-074 | 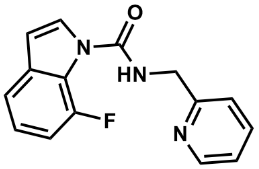 | 269.28 |
| MXC-075 | 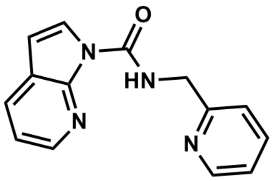 | 252.28 |
| MXC-076 | 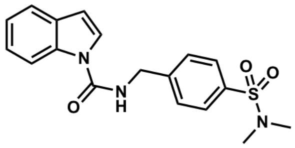 | 357.43 |
| MXC-077 | 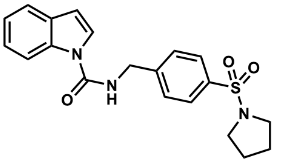 | 383.47 |

| | | |
|---|---|---|
| MXC-078 | 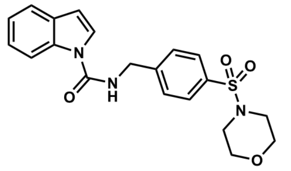 | 399.47 |
| MXC-079 | 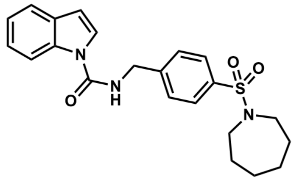 | 411.52 |
| MXC-080 | 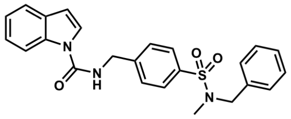 | 433.53 |
| MXC-081 | 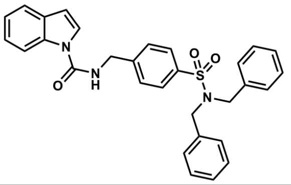 | 509.21 |
| MXC-082 | 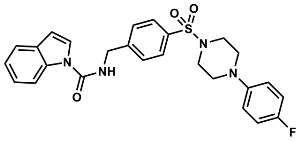 | 492.57 |
| MXC-083 | 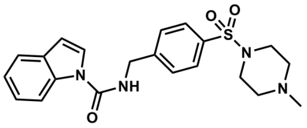 | 412.51 |
| MXC-084 | 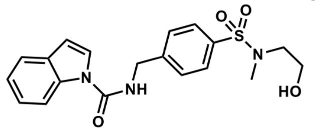 | 387.45 |
| MXC-085 | 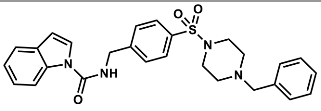 | 488.61 |
| MXC-086 | 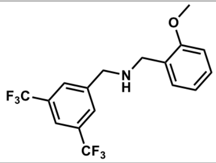 | 363.3 |
| MXC-087 | 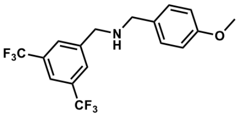 | 363.3 |
| MXC-088 | 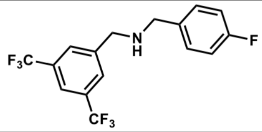 | 351.27 |

| | | |
|---|---|---|
| MXC-089 | 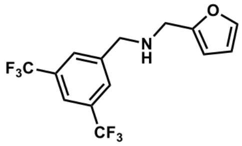 | 323.24 |
| MXC-090 | 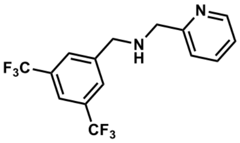 | 334.27 |
| MXC-091 | 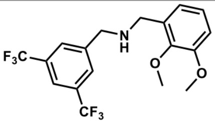 | 393.33 |
| MXC-092 | 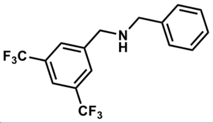 | 333.28 |
| MXC-093 | 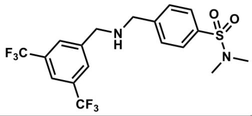 | 440.4 |
| MXC-094 | 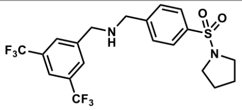 | 466.44 |
| MXC-095 | 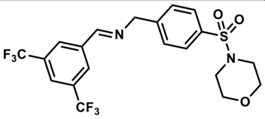 | 480.43 |
| MXC-096 | 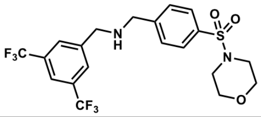 | 482.44 |
| MXC-097 | 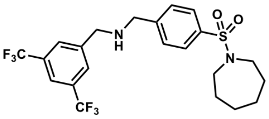 | 494.5 |
| MXC-098 | 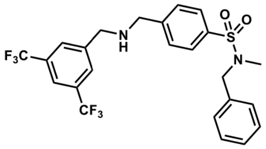 | 516.5 |
| MXC-099 | 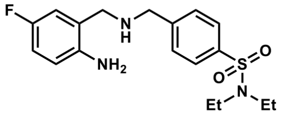 | 365.47 |
| MXC-100 | 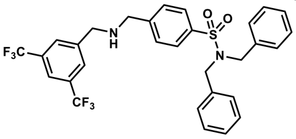 | 592.6 |
| MXC-101 | 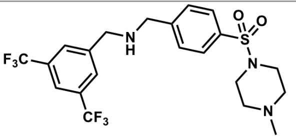 | 495.48 |

| | | |
|---|---|---|
| MXC-102 | 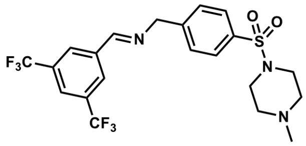 | 493.47 |
| MXC-103 | 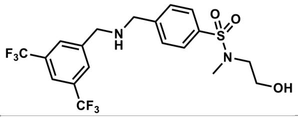 | 470.43 |
| MXC-104 | 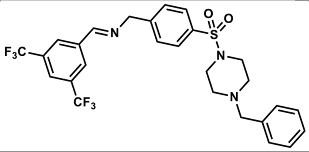 | 569.57 |
| MXC-105 | 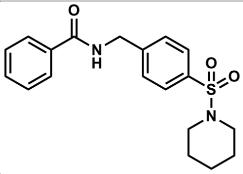 | 358.46 |
| MXC-106 | 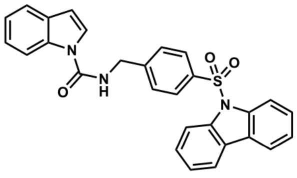 | 479.55 |
| MXC-107 | 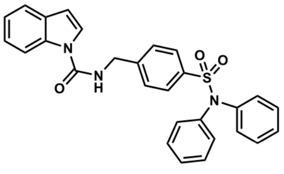 | 481.57 |
| MXC-108 | 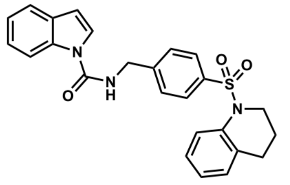 | 445.54 |
| MXC-109 | 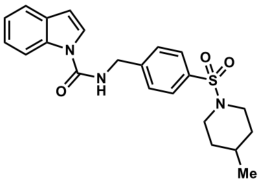 | 411.52 |
| MXC-110 | 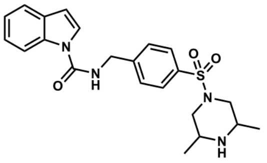 | 426.54 |
| MXC-111 | 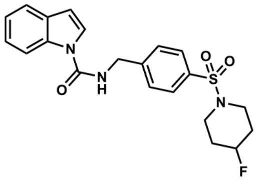 | 415.48 |

| | | |
|---|---|---|
| MXC-112 | 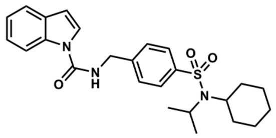 | 453.6 |
| MXC-113 | 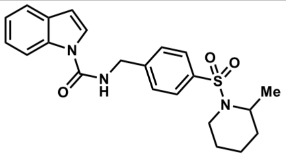 | 411.52 |
| MXC-114 | 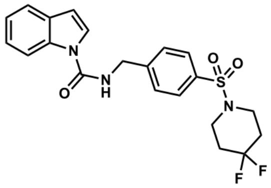 | 433.47 |

**Supplementary Table 3. Confidence intervals for stem cell frequency (%) corresponding to Figure 2.**

| Cell lines | Groups | Lower | Estimate | Upper |
|---|---|---|---|---|
| HK-374 | DMSO | 15.47987616 | 20.92050209 | 28.01120448 |
| | TMZ_1 μM | 16.75041876 | 22.62443439 | 30.12048193 |
| | TMZ_3 μM | 17.85714286 | 24.09638554 | 32.05128205 |
| | TMZ_6 μM | 10.90512541 | 14.83679525 | 20 |
| | TMZ_9 μM | 6.180469716 | 8.445945946 | 11.48105626 |
| | MXC-017_1 μM | 14.36781609 | 19.45525292 | 26.04166667 |
| | MXC-017_5 μM | 10.11122346 | 13.75515818 | 18.58736059 |
| | MXC-017_10 μM | 8.156606852 | 11.12347052 | 15.08295626 |
| | 4 Gy + DMSO | 3.875968992 | 5.327650506 | 7.299270073 |
| | 4 Gy + TMZ_1 μM | 6.447453256 | 8.865248227 | 12.12121212 |
| | 4 Gy + TMZ_3 μM | 4.504504505 | 6.191950464 | 8.481764207 |
| | 4 Gy + TMZ_6 μM | 4.030632809 | 5.540166205 | 7.593014427 |
| | 4 Gy + TMZ_9 μM | 3.374957813 | 4.638218924 | 6.361323155 |
| | 4 Gy + MXC-017_1 μM | 3.399048266 | 4.670714619 | 6.402048656 |
| | 4 Gy + MXC-017_5 μM | 3.18877551 | 4.382120947 | 6.009615385 |
| | 4 Gy + MXC-017_10 μM | 2.084636231 | 2.865329513 | 3.93236335 |
| HK-217 | DMSO | 35.33568905 | 48.7804878 | 64.1025641 |
| | TMZ_1 μM | 34.12969283 | 46.94835681 | 62.11180124 |
| | TMZ_3 μM | 21.73913043 | 30.58103976 | 41.84100418 |
| | TMZ_6 μM | 18.41620626 | 25.97402597 | 35.97122302 |
| | TMZ_9 μM | 9.389671362 | 13.42281879 | 18.97533207 |
| | MXC-017_1 μM | 24.75247525 | 34.72222222 | 47.16981132 |
| | MXC-017_5 μM | 17.54385965 | 24.81389578 | 34.36426117 |
| | MXC-017_10 μM | 9.165902841 | 13.1061599 | 18.51851852 |
| | 4 Gy + DMSO | 9.532888465 | 13.77410468 | 19.68503937 |
| | 4 Gy + TMZ_1 μM | 7.434944238 | 10.74113856 | 15.38461538 |
| | 4 Gy + TMZ_3 μM | 6.553079948 | 9.460737938 | 13.56852103 |
| | 4 Gy + TMZ_6 μM | 9.624639076 | 13.88888889 | 19.84126984 |
| | 4 Gy + TMZ_9 μM | 5.685048323 | 8.210180624 | 11.77856302 |
| | 4 Gy + MXC-017_1 μM | 7.961783439 | 11.49425287 | 16.44736842 |
| | 4 Gy + MXC-017_5 μM | 4.142502071 | 5.980861244 | 8.598452279 |
| | 4 Gy + MXC-017_10 μM | 2.346316283 | 3.387533875 | 4.880429478 |
| HK-308 | DMSO | 12.13592233 | 16.47446458 | 22.172949 |
| | TMZ_1 μM | 14.10437236 | 19.12045889 | 25.57544757 |
| | TMZ_3 μM | 15.45595054 | 20.92050209 | 27.93296089 |
| | TMZ_6 μM | 13.17523057 | 17.85714286 | 23.98081535 |
| | TMZ_9 μM | 15.43209877 | 20.87682672 | 27.93296089 |
| | MXC-017_1 μM | 13.24503311 | 17.95332136 | 24.09638554 |
| | MXC-017_5 μM | 16.28664495 | 21.97802198 | 29.3255132 |
| | MXC-017_10 μM | 7.674597084 | 10.47120419 | 14.20454545 |
| | 4 Gy + DMSO | 6.199628022 | 8.517887564 | 11.64144354 |
| | 4 Gy + TMZ_1 μM | 6.082725061 | 8.361204013 | 11.4416476 |

| | | | | |
|---|---|---|---|---|
| | 4 Gy + TMZ_3 μM | 4.543389368 | 6.242197253 | 8.554319932 |
| | 4 Gy + TMZ_6 μM | 3.231017771 | 4.440497336 | 6.090133983 |
| | 4 Gy + TMZ_9 μM | 4.42282176 | 6.079027356 | 8.326394671 |
| | 4 Gy + MXC-017_1 μM | 5.927682276 | 8.143322476 | 11.14827202 |
| | 4 Gy + MXC-017_5 μM | 6.497725796 | 8.936550492 | 12.21001221 |
| | 4 Gy + MXC-017_10 μM | 4.380201489 | 6.020469597 | 8.244023083 |
| HK-390 | DMSO | 11.58749 | 16.10306 | 22.37136 |
| | TMZ_1 μM | 7.575758 | 10.48218 | 14.51379 |
| | TMZ_3 μM | 7.390983 | 10.23541 | 14.16431 |
| | TMZ_6 μM | 5.571031 | 7.698229 | 10.6383 |
| | TMZ_9 μM | 6.002401 | 8.298755 | 11.46789 |
| | MXC-017_1 μM | 11.65501 | 16.20746 | 22.52252 |
| | MXC-017_5 μM | 7.763975 | 10.75269 | 14.88095 |
| | MXC-017_10 μM | 4.159734 | 5.740528 | 7.92393 |
| | 4 Gy + DMSO | 6.963788 | 9.699321 | 13.51351 |
| | 4 Gy + TMZ_1 μM | 4.065041 | 5.630631 | 7.800312 |
| | 4 Gy + TMZ_3 μM | 3.606203 | 4.99002 | 6.906077 |
| | 4 Gy + TMZ_6 μM | 3.937008 | 5.452563 | 7.54717 |
| | 4 Gy + TMZ_9 μM | 2.960332 | 4.091653 | 5.652911 |
| | 4 Gy + MXC-017_1 μM | 4.524887 | 6.273526 | 8.695652 |
| | 4 Gy + MXC-017_5 μM | 3.43879 | 4.757374 | 6.583278 |
| | 4 Gy + MXC-017_10 μM | 1.373061 | 1.892148 | 2.606882 |
| HK-146 | DMSO | 12.53133 | 17.4216 | 24.27184 |
| | TMZ_1 μM | 9.661836 | 13.40483 | 18.58736 |
| | TMZ_3 μM | 9.337068 | 12.93661 | 17.95332 |
| | TMZ_6 μM | 6.72043 | 9.29368 | 12.85347 |
| | TMZ_9 μM | 8.183306 | 11.33787 | 15.69859 |
| | MXC-017_1 μM | 13.45895 | 18.72659 | 26.10966 |
| | MXC-017_5 μM | 9.90099 | 13.73626 | 19.04762 |
| | MXC-017_10 μM | 5.47046 | 7.558579 | 10.43841 |
| | 4 Gy + DMSO | 5.482456 | 7.616146 | 10.58201 |
| | 4 Gy + TMZ_1 μM | 4.490346 | 6.22665 | 8.628128 |
| | 4 Gy + TMZ_3 μM | 4.152824 | 5.75374 | 7.968127 |
| | 4 Gy + TMZ_6 μM | 3.741115 | 5.178664 | 7.168459 |
| | 4 Gy + TMZ_9 μM | 3.621876 | 5.012531 | 6.934813 |
| | 4 Gy + MXC-017_1 μM | 4.882813 | 6.770481 | 9.398496 |
| | 4 Gy + MXC-017_5 μM | 2.980626 | 4.120313 | 5.694761 |
| | 4 Gy + MXC-017_10 μM | 2.328289 | 3.213368 | 4.436557 |
| HK-345 | DMSO | 10.11122 | 14.0647 | 19.53125 |
| | TMZ_1 μM | 6.858711 | 9.487666 | 13.12336 |
| | TMZ_3 μM | 7.621951 | 10.54852 | 14.59854 |
| | TMZ_6 μM | 5.144033 | 7.107321 | 9.813543 |
| | TMZ_9 μM | 4.081633 | 5.633803 | 7.770008 |
| | MXC-017_1 μM | 7.598784 | 10.51525 | 14.55604 |
| | MXC-017_5 μM | 6.501951 | 8.992806 | 12.43781 |
| | MXC-017_10 μM | 4.295533 | 5.927682 | 8.183306 |

| | | | |
|---|---|---|---|
| 4 Gy + DMSO | 5.265929 | 7.215007 | 10.15228 |
| 4 Gy + TMZ_1 μM | 3.746722 | 5.186722 | 7.178751 |
| 4 Gy + TMZ_3 μM | 3.081664 | 4.260758 | 5.889282 |
| 4 Gy + TMZ_6 μM | 3.76506 | 5.211047 | 7.215007 |
| 4 Gy + TMZ_9 μM | 3.093102 | 4.275331 | 5.910165 |
| 4 Gy + MXC-017_1 μM | 4.520796 | 6.265664 | 8.688097 |
| 4 Gy + MXC-017_5 μM | 3.355705 | 4.642526 | 6.418485 |
| 4 Gy + MXC-017_10 μM | 2.169668 | 2.994012 | 4.132231 |

**Supplementary Table 4. Pairwise tests for differences in stem cell frequencies corresponding to Figure 2.**

| Cell lines | Group 1 | Group 2 | Chisq | DF | Pr(>Chisq) |
|---|---|---|---|---|---|
| HK-374 | TMZ_1 μM | DMSO | 0.132 | 1 | 0.716 |
| | TMZ_3 μM | DMSO | 0.429 | 1 | 0.512 |
| | TMZ_6 μM | DMSO | 2.71 | 1 | 0.0998 |
| | TMZ_9 μM | DMSO | 17.5 | 1 | 2.87e-05 |
| | MXC-017_1 μM | DMSO | 0.117 | 1 | 0.733 |
| | MXC-017_5 μM | DMSO | 4.03 | 1 | 0.0448 |
| | MXC-017_10 μM | DMSO | 8.98 | 1 | 0.00273 |
| | 4 Gy + DMSO | DMSO | 36.8 | 1 | 1.28e-09 |
| | 4 Gy + TMZ_1 μM | 4 Gy + DMSO | 4.78 | 1 | 0.0288 |
| | 4 Gy + TMZ_3 μM | 4 Gy + DMSO | 0.443 | 1 | 0.505 |
| | 4 Gy + TMZ_6 μM | 4 Gy + DMSO | 0.0282 | 1 | 0.867 |
| | 4 Gy + TMZ_9 μM | 4 Gy + DMSO | 0.347 | 1 | 0.556 |
| | 4 Gy + MXC-017_1 μM | 4 Gy + DMSO | 0.312 | 1 | 0.577 |
| | 4 Gy + MXC-017_5 μM | 4 Gy + DMSO | 0.693 | 1 | 0.405 |
| | 4 Gy + MXC-017_10 μM | 4 Gy + DMSO | 7.63 | 1 | 0.00574 |
| HK-217 | TMZ_1 μM | DMSO | 0.0253 | 1 | 0.874 |
| | TMZ_3 μM | DMSO | 4.01 | 1 | 0.0451 |
| | TMZ_6 μM | DMSO | 6.82 | 1 | 0/00903 |
| | TMZ_9 μM | DMSO | 27.5 | 1 | 1.54e-07 |
| | MXC-017_1 μM | DMSO | 2.13 | 1 | 0.144 |
| | MXC-017_5 μM | DMSO | 7.99 | 1 | 0.00471 |
| | MXC-017_10 μM | DMSO | 27.6 | 1 | 1.52e-07 |
| | 4 Gy + DMSO | DMSO | 24.7 | 1 | 6.61e-07 |
| | 4 Gy + TMZ_1 μM | 4 Gy + DMSO | 0.822 | 1 | 0.364 |
| | 4 Gy + TMZ_3 μM | 4 Gy + DMSO | 1.87 | 1 | 0.171 |
| | 4 Gy + TMZ_6 μM | 4 Gy + DMSO | 0.00125 | 1 | 0.972 |
| | 4 Gy + TMZ_9 μM | 4 Gy + DMSO | 3.48 | 1 | 0.0619 |
| | 4 Gy + MXC-017_1 μM | 4 Gy + DMSO | 0.433 | 1 | 0.511 |
| | 4 Gy + MXC-017_5 μM | 4 Gy + DMSO | 9.3 | 1 | 0.0023 |
| | 4 Gy + MXC-017_10 μM | 4 Gy + DMSO | 26.9 | 1 | 2.1e-07 |
| HK-308 | TMZ_1 μM | DMSO | 0.496 | 1 | 0.481 |
| | TMZ_3 μM | DMSO | 1.22 | 1 | 0.27 |
| | TMZ_6 μM | DMSO | 0.143 | 1 | 0.706 |
| | TMZ_9 μM | DMSO | 1.26 | 1 | 0.262 |
| | MXC-017_1 μM | DMSO | 0.164 | 1 | 0.686 |
| | MXC-017_5 μM | DMSO | 1.94 | 1 | 0.163 |
| | MXC-017_10 μM | DMSO | 4.5 | 1 | 0.0338 |
| | 4 Gy + DMSO | DMSO | 8.71 | 1 | 0.00316 |
| | 4 Gy + TMZ_1 μM | 4 Gy + DMSO | 0.00599 | 1 | 0.938 |
| | 4 Gy + TMZ_3 μM | 4 Gy + DMSO | 1.66 | 1 | 0.197 |
| | 4 Gy + TMZ_6 μM | 4 Gy + DMSO | 7.23 | 1 | 0.00715 |

| | | | | | |
|---|---|---|---|---|---|
| | 4 Gy + TMZ_9 µM | 4 Gy + DMSO | 1.95 | 1 | 0.163 |
| | 4 Gy + MXC-017_1 µM | 4 Gy + DMSO | 0.0345 | 1 | 0.853 |
| | 4 Gy + MXC-017_5 µM | 4 Gy + DMSO | 0.0397 | 1 | 0.842 |
| | 4 Gy + MXC-017_10 µM | 4 Gy + DMSO | 2.11 | 1 | 0.146 |
| HK-390 | TMZ_1 µM | DMSO | 3.06 | 1 | 0.0804 |
| | TMZ_3 µM | DMSO | 3.44 | 1 | 0.0635 |
| | TMZ_6 µM | DMSO | 9.2 | 1 | 0.00242 |
| | TMZ_9 µM | DMSO | 7.21 | 1 | 0.00723 |
| | MXC-017_1 µM | DMSO | 0.000548 | 1 | 0.981 |
| | MXC-017_5 µM | DMSO | 2.68 | 1 | 0.101 |
| | MXC-017_10 µM | DMSO | 18.2 | 1 | 1.96e-05 |
| | 4 Gy + DMSO | DMSO | 4.19 | 1 | 0.0407 |
| | 4 Gy + TMZ_1 µM | 4 Gy + DMSO | 5.03 | 1 | 0.0249 |
| | 4 Gy + TMZ_3 µM | 4 Gy + DMSO | 7.73 | 1 | 0.00542 |
| | 4 Gy + TMZ_6 µM | 4 Gy + DMSO | 5.62 | 1 | 0.0178 |
| | 4 Gy + TMZ_9 µM | 4 Gy + DMSO | 12.5 | 1 | 0.000408 |
| | 4 Gy + MXC-017_1 µM | 4 Gy + DMSO | 3.27 | 1 | 0.0707 |
| | 4 Gy + MXC-017_5 µM | 4 Gy + DMSO | 8.6 | 1 | 0.00336 |
| | 4 Gy + MXC-017_10 µM | 4 Gy + DMSO | 50.3 | 1 | 1.31e-12 |
| HK-146 | TMZ_1 µM | DMSO | 1.14 | 1 | 0.286 |
| | TMZ_3 µM | DMSO | 1.48 | 1 | 0.223 |
| | TMZ_6 µM | DMSO | 6.56 | 1 | 0.0104 |
| | TMZ_9 µM | DMSO | 3.04 | 1 | 0.0811 |
| | MXC-017_1 µM | DMSO | 4.24 | 1 | 0.0395 |
| | MXC-017_5 µM | DMSO | 0.943 | 1 | 0.331 |
| | MXC-017_10 µM | DMSO | 11.3 | 1 | 0.000765 |
| | 4 Gy + DMSO | DMSO | 11.1 | 1 | 0.000851 |
| | 4 Gy + TMZ_1 µM | 4 Gy + DMSO | 0.679 | 1 | 0.41 |
| | 4 Gy + TMZ_3 µM | 4 Gy + DMSO | 1.28 | 1 | 0.257 |
| | 4 Gy + TMZ_6 µM | 4 Gy + DMSO | 2.45 | 1 | 0.118 |
| | 4 Gy + TMZ_9 µM | 4 Gy + DMSO | 2.87 | 1 | 0.0904 |
| | 4 Gy + MXC-017_1 µM | 4 Gy + DMSO | 0.226 | 1 | 0.634 |
| | 4 Gy + MXC-017_5 µM | 4 Gy + DMSO | 6.36 | 1 | 0.0117 |
| | 4 Gy + MXC-017_10 µM | 4 Gy + DMSO | 12 | 1 | 0.00053 |
| HK-345 | TMZ_1 µM | DMSO | 2.64 | 1 | 0.104 |
| | TMZ_3 µM | DMSO | 1.37 | 1 | 0.241 |
| | TMZ_6 µM | DMSO | 7.88 | 1 | 0.005 |
| | TMZ_9 µM | DMSO | 14.3 | 1 | 0.00016 |
| | MXC-017_1 µM | DMSO | 1.43 | 1 | 0.232 |
| | MXC-017_5 µM | DMSO | 3.43 | 1 | 0.0642 |
| | MXC-017_10 µM | DMSO | 12.4 | 1 | 0.000426 |
| | 4 Gy + DMSO | DMSO | 7.05 | 1 | 0.00792 |
| | 4 Gy + TMZ_1 µM | 4 Gy + DMSO | 1.97 | 1 | 0.161 |
| | 4 Gy + TMZ_3 µM | 4 Gy + DMSO | 4.92 | 1 | 0.0265 |
| | 4 Gy + TMZ_6 µM | 4 Gy + DMSO | 1.87 | 1 | 0.171 |
| | 4 Gy + TMZ_9 µM | 4 Gy + DMSO | 4.67 | 1 | 0.0306 |

| | | | | | |
|---|---|---|---|---|---|
| | 4 Gy + MXC-017_1 µM | 4 Gy + DMSO | 0.387 | 1 | 0.534 |
| | 4 Gy + MXC-017_5 µM | 4 Gy + DMSO | 3.36 | 1 | 0.0668 |
| | 4 Gy + MXC-017_10 µM | 4 Gy + DMSO | 13.2 | 1 | 0.000273 |

**Supplementary Table 5. Confidence intervals for stem cell frequency (%) corresponding to Figure 3.**

| Groups | Lower | Estimate | Upper |
|---|---|---|---|
| Corn oil | 0.82644628 | 1.11982083 | 1.51515152 |
| MXC-017 | 0.37593985 | 0.47528517 | 0.5988024 |
| 4 Gy + corn oil | 0.19762846 | 0.32414911 | 0.36101083 |
| 4 Gy + MXC-017 | 6.24649e-003 | 0 | 0 |

**Supplementary Table 6. Pairwise tests for differences in stem cell frequencies corresponding to Figure 3.**

| Group 1 | Group 2 | Chisq | DF | Pr(>Chisq) |
|---|---|---|---|---|
| MXC-017 | Corn oil | 20.8 | 1 | 5.18e-06 |
| 4 Gy + corn oil | Corn oil | 9.23 | 1 | 0.00238 |
| 4 Gy + MXC-017 | Corn oil | 243 | 1 | 8.34e-55 |
| 4 Gy + MXC-017 | 4 Gy + corn oil | 127 | 1 | 2.32e-29 |

**Supplementary Table 7. Cell Type Composition and Relative Proportions in HK-217 i*n vitro* scRNA-seq Dataset**

| Sample | Cell type | Count | Proportion (%) |
|---|---|---|---|
| Control | Astrocyte | 14 | 0.19 |
| | Cycling | 3049 | 40.53 |
| | MES | 19 | 0.25 |
| | Microglia | 6 | 0.08 |
| | NPC | 1 | 0.01 |
| | OPC | 2815 | 37.42 |
| | Oligo | 601 | 8.00 |
| | RG | 392 | 5.21 |
| | Vascular | 621 | 8.26 |
| | oRG | 4 | 0.05 |
| MXC-017 | Cycling | 1842 | 41.84 |
| | MES | 13 | 0.30 |
| | Microglia | 7 | 0.16 |
| | NPC | 5 | 0.11 |
| | Neuronal | 2 | 0.05 |
| | OPC | 1571 | 35.68 |
| | Oligo | 480 | 10.90 |
| | RG | 315 | 7.15 |
| | Vascular | 161 | 3.66 |
| | oRG | 7 | 0.16 |
| RT | Astrocyte | 62 | 0.44 |
| | Cycling | 7466 | 52.94 |
| | MES | 55 | 0.39 |
| | Microglia | 23 | 0.16 |
| | NPC | 18 | 0.13 |
| | Neuronal | 1 | 0.01 |
| | OPC | 3480 | 24.67 |
| | Oligo | 1487 | 10.54 |
| | RG | 1302 | 9.23 |
| | Vascular | 201 | 1.43 |
| | oRG | 9 | 0.06 |
| RT+MXC-017 | Astrocyte | 30 | 0.23 |
| | Cycling | 5870 | 45.01 |
| | MES | 32 | 0.25 |
| | Microglia | 28 | 0.21 |
| | NPC | 18 | 0.14 |
| | Neuronal | 7 | 0.05 |
| | OPC | 3886 | 29.80 |
| | Oligo | 1333 | 10.22 |
| | RG | 1596 | 12.24 |
| | Vascular | 217 | 1.66 |
| | oRG | 25 | 0.19 |

**Supplementary Table 8. Cell Type Composition and Relative Proportions in HK-244 i*n vitro* scRNA-seq Dataset**

| Sample | Cell type | Count | Proportion (%) |
|---|---|---|---|
| Control | Astrocyte | 4 | 0.05 |
| | Cycling | 6874 | 86.41 |
| | MES Progenitor | 1 | 0.01 |
| | Microglia | 5 | 0.06 |
| | Neuronal | 138 | 1.73 |
| | OPC | 457 | 5.74 |
| | RG | 65 | 0.82 |
| | Transition State | 36 | 0.45 |
| | Undefined Progenitor | 17 | 0.21 |
| | Vascular | 32 | 0.40 |
| | oRG | 326 | 4.10 |
| MXC-017 | Astrocyte | 18 | 0.53 |
| | Cycling | 2343 | 69.57 |
| | MES | 2 | 0.06 |
| | MES Progenitor | 1 | 0.03 |
| | Microglia | 179 | 5.31 |
| | NPC | 15 | 0.45 |
| | Neuronal | 256 | 7.60 |
| | OPC | 168 | 5.00 |
| | RG | 43 | 1.28 |
| | Transition State | 23 | 0.68 |
| | Vascular | 57 | 1.70 |
| | oRG | 263 | 7.81 |
| RT | Astrocyte | 16 | 0.33 |
| | Cycling | 3739 | 77.31 |
| | MES | 1 | 0.02 |
| | MES Progenitor | 9 | 0.19 |
| | Microglia | 217 | 4.49 |
| | NPC | 9 | 0.19 |
| | Neuronal | 254 | 5.25 |
| | OPC | 176 | 3.64 |
| | RG | 42 | 0.87 |
| | Transition State | 8 | 0.17 |
| | Undefined Progenitor | 1 | 0.02 |
| | Vascular | 69 | 1.43 |
| | oRG | 295 | 6.10 |
| RT+MXC-017 | Astrocyte | 16 | 0.36 |
| | Cycling | 3362 | 76.03 |
| | MES | 1 | 0.02 |
| | MES Progenitor | 5 | 0.11 |
| | Microglia | 212 | 4.79 |
| | NPC | 9 | 0.20 |

| | | | |
|---|---|---|---|
| | Neuronal | 249 | 5.63 |
| | OPC | 170 | 3.84 |
| | RG | 31 | 0.70 |
| | Transition State | 8 | 0.18 |
| | Undefined Progenitor | 1 | 0.02 |
| | Vascular | 68 | 1.54 |
| | oRG | 290 | 6.56 |

**Supplementary Table 9. Cell Type Composition and Relative Proportions in HK-374 i*n vitro* scRNA-seq Dataset**

| Sample | Cell type | Count | Proportion (%) |
|---|---|---|---|
| Control | Astrocyte | 64 | 0.73 |
| | Cycling | 2516 | 28.68 |
| | MES | 4456 | 50.80 |
| | Microglia | 10 | 0.11 |
| | Mixed_Vascular | 58 | 0.66 |
| | RG | 408 | 4.65 |
| | Vascular | 1226 | 13.98 |
| | oRG | 34 | 0.39 |
| MXC-017 | Astrocyte | 10 | 0.15 |
| | Cycling | 2974 | 44.47 |
| | MES | 3253 | 48.65 |
| | Microglia | 1 | 0.01 |
| | Mixed_Vascular | 12 | 0.18 |
| | RG | 164 | 2.45 |
| | Undefined Progenitor | 4 | 0.06 |
| | Vascular | 262 | 3.92 |
| | oRG | 7 | 0.10 |
| RT | Cycling | 263 | 6.37 |
| | MES | 3555 | 86.16 |
| | Microglia | 3 | 0.07 |
| | Mixed_Vascular | 14 | 0.34 |
| | RG | 214 | 5.19 |
| | Undefined Progenitor | 1 | 0.02 |
| | Vascular | 73 | 1.77 |
| | oRG | 3 | 0.07 |
| RT+MXC-017 | Cycling | 334 | 6.87 |
| | MES | 4011 | 82.45 |
| | Microglia | 8 | 0.16 |
| | Mixed_Vascular | 20 | 0.41 |
| | RG | 303 | 6.23 |
| | Vascular | 171 | 3.51 |
| | oRG | 18 | 0.37 |

**Supplementary Table 10. Cell Type Composition and Relative Proportions in HK-390 i*n vitro* scRNA-seq Dataset**

| Sample | Cell type | Count | Proportion (%) |
|---|---|---|---|
| Control | Cycling | 8376 | 82.70 |
| | MES Progenitor | 41 | 0.40 |
| | Microglia | 81 | 0.80 |
| | NPC | 1 | 0.01 |
| | Neuronal | 1 | 0.01 |
| | RG | 291 | 2.87 |
| | Vascular | 9 | 0.09 |
| | oRG | 1328 | 13.11 |
| MXC-017 | Cycling | 8410 | 87.38 |
| | MES Progenitor | 58 | 0.60 |
| | Microglia | 13 | 0.14 |
| | RG | 273 | 2.84 |
| | Vascular | 2 | 0.02 |
| | oRG | 869 | 9.03 |
| RT | Cycling | 4615 | 80.21 |
| | MES | 1 | 0.01 |
| | MES Progenitor | 103 | 1.80 |
| | Microglia | 61 | 1.06 |
| | RG | 414 | 7.19 |
| | Vascular | 12 | 0.21 |
| | oRG | 548 | 9.52 |
| RT+MXC-017 | Cycling | 6503 | 81.17 |
| | MES | 2 | 0.02 |
| | MES Progenitor | 122 | 1.52 |
| | Microglia | 47 | 0.59 |
| | RG | 579 | 7.23 |
| | Vascular | 8 | 0.10 |
| | oRG | 751 | 9.37 |

**Supplementary Table 11. Cell Type Composition and Relative Proportions in HK-374 *in vivo* scRNA-seq Dataset**

| Sample | Cell type | Count | Proportion (%) |
|---|---|---|---|
| Control | Astrocyte | 2 | 0.03 |
| | Cycling | 654 | 11.26 |
| | MES | 3323 | 57.22 |
| | Microglia | 2 | 0.03 |
| | Mixed_Vascular | 1770 | 30.48 |
| | Neuronal | 2 | 0.03 |
| | Oligo | 1 | 0.02 |
| | RG | 24 | 0.41 |
| | oRG | 29 | 0.50 |
| RT | Cycling | 291 | 8.36 |
| | MES | 2129 | 61.18 |
| | Microglia | 4 | 0.11 |
| | Mixed_Vascular | 1019 | 29.28 |
| | Neuronal | 9 | 0.26 |
| | RG | 8 | 0.23 |
| | oRG | 20 | 0.57 |
| RT+MXC-017_1 | Astrocyte | 12 | 1.18 |
| | Cycling | 83 | 8.14 |
| | MES | 573 | 56.18 |
| | Microglia | 3 | 0.29 |
| | Mixed_Vascular | 314 | 30.78 |
| | Neuronal | 3 | 0.29 |
| | Oligo | 4 | 0.39 |
| | RG | 6 | 0.59 |
| | oRG | 22 | 2.16 |
| RT+MXC-017_2 | Astrocyte | 3 | 0.50 |
| | Cycling | 40 | 6.67 |
| | MES | 335 | 55.83 |
| | Mixed_Vascular | 152 | 25.33 |
| | Neuronal | 55 | 9.17 |
| | Oligo | 1 | 0.17 |
| | RG | 3 | 0.50 |
| | oRG | 11 | 1.83 |
